# The influence of circadian rhythms on CD8^+^ T cell activation upon vaccination: a mathematical modeling perspective

**DOI:** 10.1101/2023.09.28.559961

**Authors:** Nasri Balit, Nicolas Cermakian, Anmar Khadra

**Affiliations:** Department of Physiology, McGill University, Montreal, Quebec, Canada; Douglas Research Center, Department of Psychiatry, McGill University, Montreal, Quebec, Canada

**Keywords:** CD8^+^ T cells, Adaptive immunity, Vaccination dynamics, Circadian rhythm, Mathematical modeling, Bifurcation analysis, Bistable switch, PRCC, Population level bistable switch

## Abstract

Circadian rhythms have been implicated in the modulation of many physiological processes, including those associated with the immune system. For example, these rhythms influence CD8^+^ T cell responses within the adaptive immune system. The mechanism underlying this immune-circadian interaction, however, remains unclear, particularly in the context of vaccination. Here, we devise a molecularly-explicit gene regulatory network model of early signaling in the näıve CD8^+^ T cell activation pathway, comprised of three axes (or subsystems) labeled ZAP70, LAT and CD28, to elucidate the molecular details of this immune-circadian mechanism and its relation to vaccination. This is done by coupling the model to a periodic forcing function to identify the molecular players targeted by circadian rhythms, and analyzing how these rhythms subsequently affect CD8^+^ T cell activation under differing levels of T cell receptor (TCR) phosphorylation, which we designate as vaccine load. By performing both bifurcation and parameter sensitivity analyses on the model at the single cell and population levels, we find that applying periodic forcing on molecular targets within the ZAP70 axis is sufficient to create a day-night discrepancy in CD8^+^ T cell activation in a manner that is dependent on the bistable switch inherent in CD8^+^ T cell early signaling. We also demonstrate that the resulting CD8^+^ T cell activation is dependent on the strength of the periodic coupling as well as on the level of TCR phosphorylation. Our results show that this day-night discrepancy is not transmitted to certain downstream molecules within the LAT subsystem, such as mTORC1, suggesting a secondary, independent circadian regulation on that protein complex. We also corroborate experimental results by showing that the circadian regulation of CD8^+^ T cell primarily acts at a baseline, pre-vaccination state, playing a facilitating role in priming CD8^+^ T cells to vaccine inputs according to time of day. By applying a population level analysis using bifurcation theory and by including several hypothesized molecular targets of this circadian rhythm, we further demonstrate an increased variability between CD8^+^ T cells (due to heterogeneity) induced by its circadian regulation, which may allow a population of CD8^+^ T cells to activate at a lower vaccine load, improving its sensitivity. This modeling study thus provides insights into the immune targets of the circadian clock, and proposes an interaction between vaccine load and the influence of circadian rhythms on CD8^+^ T cell activation.

**Highlights:**

- Potential targets of circadian rhythms within the ZAP70 signaling pathway were identified.
- The level of vaccine load to a CD8^+^ T cell was shown to be crucial in dictating a circadian rhythm’s influence on its signaling response.
- The ’priming’ effect of a circadian rhythm on CD8^+^ T cell activation upon vaccination was demonstrated.
- mTORC1 and its immediate upstream signaling molecules were shown to be regulated by circadian rhythms through independent mechanisms.
- A heterogeneity in CD8^+^ T cells, induced by its circadian regulation, may influence their sensitivity to vaccination.

## 1. Introduction

Circadian rhythms, 24 h cycles that modulates many of the body’s internal processes (Portaluppi et al., 2012; de Goede et al., 2018), have been linked to the immune system from as far back as the 1960s (Halberg et al., 1960; Esquifino et al., 1996; Kawate et al., 1981; Young et al., 1995). This modulation is thought to be regulated by the presence of molecular clocks, made of clock genes, within cells of the immune system (Houtman et al., 2005; An et al., 2019; De Boer et al., 2001; Coombs et al., 2011; Bollinger et al., 2011; Cuesta et al., 2017; Boivin et al., 2003), including *Clock, Bmal1, Per1-3, Cry1-2*) (Dibner et al., 2010). The expression level of these clock genes is regulated by transcription-translation feedback loops that create oscillations in their expression pattern with a *∼*24 h period. These clock genes also modulate the expression of specific molecular targets within immune cells in a cyclical manner, imposing circadian rhythms on them (Labrecque and Cermakian, 2015; Jerigova et al., 2022). With evidence suggesting that circadian rhythms affect how the immune system responds to vaccination (Hazan et al., 2023; Cermakian et al., 2021; Elliott et al., 1972; Long et al., 2016; Phillips et al., 2008; de Bree et al., 2020), it becomes imperative to understand how this relation is manifested molecularly and dynamically at the single cell and population levels.

The molecular basis of the immune-circadian interaction has been previously linked to specific processes within the innate immune system (Labrecque and Cermakian, 2015; Jerigova et al., 2022). Indeed, molecular clocks have been characterized in macrophages, neutrophils, eosinophils and natural killer (NK) cells (Keller et al., 2009). For example, in macrophages, the clock gene *REV-ERBα* has been shown to affect the inflammatory function of macrophages by repressing their NF-*κ*B activation pathway (Sato et al., 2014; Wang et al., 2018), whereas in NK cells, mice with *Per2* knockdowns displayed reduced levels of proinflammatory cytokines, namely IFN-*γ* and IL-1*β*, after an LPS endotoxic shock (Liu et al., 2006). Furthermore, knockouts of *Bmal1* in myeloid cells predispose mice to developing acute and chronic inflammation pathologies by disrupting the diurnal variation in monocyte number (Nguyen et al., 2013).

Although significant progress has been made in understanding how innate immunity is influenced by circadian rhythms, the molecular basis of this immune-circadian link remains unclear in adaptive immunity, particularly in CD8^+^ effector T cell response. While circadian oscillations were observed in the RNA level of *Per2* in lymph node (Fortier et al., 2011; Nobis et al., 2019a) and circulating näıve CD8^+^ T cells (Dimitrov et al., 2009), the molecular source(s) of this daily variation remains unknown. Additionally, this circadian variation in CD8^+^ T cell activity may, in part, depend on cell-autonomous mechanisms, such as clocks within those cells (Nobis et al., 2019a; Wang et al., 2023). Through transcriptomic analysis, a recent study identified several genes within the CD8^+^ T cell activation pathway that exhibited rhythmicity in their expression in näıve CD8^+^ T cells, and subsequently demonstrated that certain molecular players within this activation pathway, such as ZAP70, AKT and mTOR, demonstrated an increased activation following a day-time vaccination, as well as an increased ability to combat an infectious challenge after a day-time vaccination (both compared to night-time vaccination) (Nobis et al., 2019b). Moreover, another study showed an increased level of ZAP70 prior to vaccination in the day-time (Fortier et al., 2011). These studies thus suggest that certain molecular players within the CD8^+^ T cell activation signaling pathway interact with circadian rhythms in a way that primes these T cells to exhibit increased activation when vaccinated during the day-time compared to night-time; the details of this interaction, however, remains not fully explored.

In this study, we generate a theoretical framework, through mathematical modeling, to further elucidate the molecular basis of the immune-circadian link in CD8^+^ T cells following a vaccine input and its underlying dynamics. We adapt an experimentally validated, CD4^+^ T cell gene regulatory model (Zheng et al., 2005; Perley et al., 2014) to simulate the immediate signaling events during näıve CD8^+^ T cell activation following TCR phosphorylation. By expanding the model to include a parameter-specific periodic forcing function, we explore the effects of a circadian rhythm on this signaling pathway to determine how it impacts CD8^+^ T cell activation postvaccination in a molecular-player-specific manner. This is first done via a single-parameter model framework, where bifurcation, sensitivity and time series analyses are performed on a selected parameter to develop a deeper understanding of the dynamics of this modulation. We then expand this approach to a multi-parameter framework to generate more physiologically relevant conclusions about this system. While prior modeling studies have explored the effect of a circadian rhythm on the immune system (Bai and Zhou, 2012; Fan and Wang, 2010; Ji et al., 2010; Wang et al., 2006), these studies were not done using T cell gene regulatory models.

Through our single cell and population level modeling, we demonstrate the crucial role played by the ZAP70 axis (or subsystem) within the immune activation pathway in how a circadian rhythm exerts its effects in modulating CD8^+^ T cell responses. By first highlighting the presence of a bistable switch in this subsystem, we show that this switch dictates the dynamics of this circadian modulation, and that the latter is dependent on both the number of phosphorylated T cell receptors (TCRs), which we term the vaccine load, presented to the cell, as well as on the magnitude of circadian coupling. Additionally, we establish that this modulation occurs prior to vaccination, suggesting a priming role for a circadian rhythm on CD8^+^ T cell signaling. Interestingly, our results also reveal that downstream molecular players, such as the mTORC1 protein complex, may be independently regulated by circadian rhythms, pointing to the existence of a complex, multi-layered circadian influence on CD8^+^ T cell signaling. Our population modeling further expand on these ideas using a multi-parameter framework to produce more physio-logical conclusions about the influence of a circadian rhythm on CD8^+^ T cell activation.

## 2. Single-cell model framework

### 2.1. Gene regulatory network model

To explore the dynamics of circadian influence on CD8^+^ T cell activation, a multi-scale gene regulatory network model of the early events of T cell signaling is considered. The network model is described by a system of ordinary differential equations (ODEs) adapted from literature. This model, first developed in Zheng et al. (2005) and later extended in Perley et al. (2014), describes the immediate signaling events following antigenic stimulation of a single CD4^+^ T cell. Because of the overlap in these early signaling events between CD8^+^ and CD4^+^ T cells (Mørch et al., 2020), we have adapted the CD4^4^ model to describe CD8^+^ T cell signaling and modified it to exclude the gene regulatory output signals described in Perley et al. (2014) since CD4^+^ and CD8^+^ näıve T cells vary in their gene outputs (Hosking et al., 2014). The model is then coupled to a rhythmic signal representing a circadian rhythm and a single rate-activation parameter is chosen as a molecular target for circadian rhythm coupling. The effects of this coupling on model dynamics are subsequently analyzed.

Briefly, the gene regulatory network model is comprised of the early signaling events following T cell receptor (TCR) phosphorylation (Fig.1). According to this framework, a phosphorylated TCR (denoted by *TCRp*) binds to a tyrosine kinase (*ZAP* 70) and triggers a cascade of interactions that ultimately result in the binding of *ZAP* 70 to activated Src family protein kinases (*SFK*). This cascade of interactions is governed by a complex network of feedback mechanisms, regulated by positive feedback loops from the various forms of *ZAP* 70 (denoted by *ZAP ^∗^, ZAPp* and *SFK^∗^*-*ZAPp*), as well as negative feedback loops from Src homology region 2 domain-containing phosphatase-1 (*SHP* 1), C-terminal Src kinase (*CSK*), and CREB-binding protein (*CBP*). The activated *ZAPp* and *SFK^∗^*-*ZAPp* then feed into a series of reactions governed by various molecular players, such *LAT, AKT* etc., which ultimately result in the production of mammalian target of rapamycin complex 1 & 2 (*mTORC*1 and *mTORC*2). Both of these protein complexes, *mTORC*1 and *mTORC*2, are important in determining T cell fate and effector function (Chi, 2012). The rate equations and parameter values for this model are shown in Appendix 6.1, and a more detailed description can be found in Zheng et al. (2005) and Perley et al. (2014). The rate equations and parameter values of this model are provided in Table 6.1.

**Figure 1:**
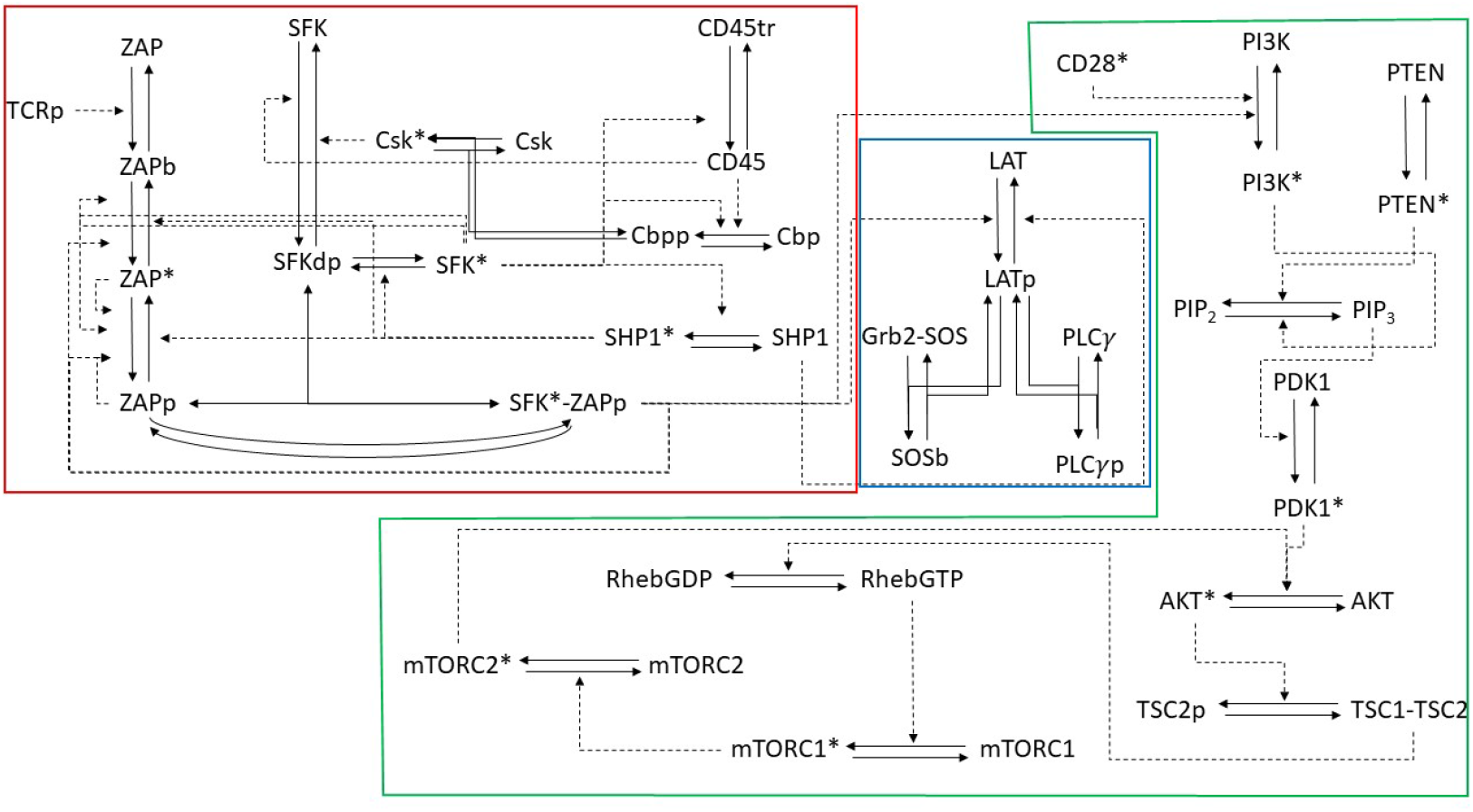
Schematic of the gene regulatory network adapted from Zheng et al. (2005) and Perley et al. (2014), and considered in our model framework to describe early signaling in CD8^+^ T cells. The model can be divided into three main subsystems: the ZAP axis (red box), LAT axis (blue box) and CD28 axis (green box). Notice how the LAT and CD28 axes are linked to the ZAP axis in a feedforward manner without any feedback, and that the LAT and CD28 axes are similarly not linked in a feedforward manner. Asterisk is used to represent the active forms of the relevant molecules.

### 2.2. The three key axes of the model

To explore the impact of a circadian rhythm on different components of the model, we divide the gene regulatory network model (Fig. 1) into three subsystems based on how they feed into each other. The first subsystem (labeled the ZAP axis and highlighted by the red box in Fig. 1) includes the reactions that involve *ZAP_active_* formation. It begins with the *TCRp* input, and ends with *SFK ∗ −ZAPp* and *SHP* 1. The second subsystem involves the signaling molecules downstream of *ZAP_active_*, which include the reactions governing LAT phosphorylation (hence the label LAT axis highlighted by the blue box in Fig. 1). These reactions similarly do not receive feedback from downstream signaling molecules. Finally, the third subsystem involves the signaling molecules included in the *CD*28 signaling pathway (hence the label CD28 axis highlighted by the green box in Fig. 1)). It starts with the two input signals: *ZAP_active_* and *CD*28, and ends with *mTORC*1*^∗^*. By exploring the effect of a circadian rhythm on the output of each one of these axes and determining how this circadian influence is filtered from one axis to the next, one can understand which axis is more sensitive to circadian-induced perturbations, helping us identify potential molecular targets of this circadian rhythm.

### 2.3. Periodic forcing by circadian rhythms

To investigate the effect(s) of a circadian rhythm on T cell early signaling events, we incorporate a parameter-specific periodic forcing function applied on a given rate constant of one of these early signaling events; that is to say, an external forcing function is imposed onto a parameter of interest within the model, *p*_0_, to produce a time-dependent variation in its value, denoted by *p_t_*. This time-dependent variation in *p*_0_ may get transmitted to downstream signals within the model, producing a time-dependent difference in its output signal. If so, this will allow us to identify which parameter(s) is/are responsible for the day-night variation in CD8^+^ T cell signaling. The equation we choose for this periodic forcing is given by

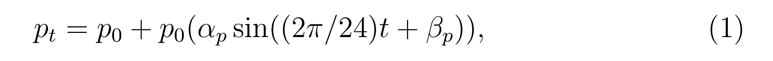

where *p*_0_ is the default (original) value of the parameter *p* that is directly affected by the periodic forcing representing a circadian rhythm, *t* is time, *α_p_* is the parameter-specific ’circadian coefficient’ that determines how strongly that parameter is impacted by the oscillation, and *β_p_* determines the parameter-specific phase shift of the oscillation.

According to Eqn. 1, the periodic forcing is defined as a sine function that is oscillating with a mean *p*_0_ and a circadian coefficient *α*, representing the magnitude of circadian rhythm coupling to a specific parameter of interest. Such description allows us to study the effect of different strengths of circadian coupling on model dynamics. The choice of a sine function to describe circadian rhythms is consistent with how it is routinely characterized in experimental literature (Elliott et al., 1972; Hickey et al., 1984; Naitoh et al., 1985).

To account for physiological limitations of how much a parameter can vary (see Appendix 6.3.2), we assume that *α_p_ ∈* [0, 0.5]. The periodic forcing defined in Eqn. 1 produces oscillations with a 24 h period and a phase shift of 0 for a positive feedback parameter and *π* for a negative feedback parameter; this reproduces the effect of a circadian rhythm, and leads to a variation in the signaling molecule associated with *p_t_*, depending on time of day. Increasing the circadian coefficient *α* causes the amplitude of *p_t_* to increase and thus the range of values of *p_t_* to expand in a parameter-specific manner.

## 3. Outcomes of the single cell model

### 3.1. Excluding the oscillatory regime

The original parameterization for the ZAP axis described by Zheng et al. (2005) is capable of producing intrinsic, yet unphysiological, oscillatory dynamics in T cell activation (see Appendix 6.2). To resolve this issue, we have adapted a Dynamics Visualisation based Parallel Coordinates (DYVIPAC) framework introduced by Nguyen et al. (2015) to identify the oscillatory regime and exclude it (see Appendix 6.3). This has allowed us to confine model’s parameter values (as well as parameter sampling performed in subsequent sections) to a connected non-oscillatory parameter regime, ensuring that oscillations observed in the model result exclusively from the periodic forcing function representing the 24-hour circadian cycle.

### 3.2. Strategy for identifying the molecular targets of the CD8^+^ T cell circadian rhythm

Identifying the parameters responsible for producing the key features of model dynamics will provide us with potential targets for producing the day-night variation in model output. By performing a dynamical systems analysis of the model, one can predict how parameter perturbations, due to the CD8^+^ T cell circadian rhythm, can modify model behaviour. To accomplish this, we make the assumption here that a circadian rhythm on CD8^+^ T cell signaling would primarily affect the rate activation kinetics of signaling molecules within the CD8^+^ T cell activation pathway. This is supported by the day-night variation in active forms of CD8^+^ T cell signaling molecules shown in Nobis et al. (Nobis et al., 2019b).

We begin first by performing parameter sweeps and bifurcation analyses within each axis of the gene regulatory network model to identify the broad signaling pathways defining the dynamics of our model. Once a ’key’ axis is found, individual parameters within the axis are further explored using both parameter sensitivity and threshold analyses to identify targets for this circadian rhythm (i.e., periodic forcing).

#### 3.2.1. The ZAP axis is pivotal in defining model dynamics

The ZAP axis is the most upstream subsystem of the model, receiving a direct *TCRp* input and outputting various forms of activated ZAP: phosphorylated ZAP (*ZAPp*), active ZAP (*ZAP ^∗^*), and active SFK-bound ZAP (*SFK^∗^*-*ZAPp*). To consolidate the various active forms of ZAP into a single indicator variable, we use the same approach as in Zheng et al. (2005) by defining a new output variable *ZAP_active_* for this axis, given by

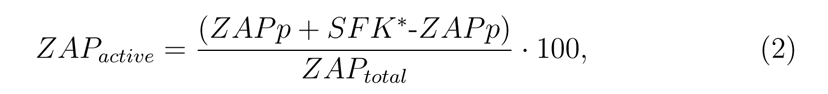

where *ZAP_active_* is the percentage of ZAP within a single T cell that has been fully phosphorylated, ZAPp and SFK*-ZAPp are the fully phosphorylated forms of ZAP70, and *ZAP_total_* denotes the conserved total quantity of ZAP. With this output variable, the dynamics of the ZAP axis are explored through parameter sweeps and bifurcation analysis. In Fig. 2(A), we show that for *TCRp* = 27000 mol, the time course of *ZAP_active_*, the output signal of the ZAP axis, changes significantly as the values of the rate activation parameters of the model are altered. For instance, as the rate activation parameter for SHP1, *k*10*f* (blue) decreases past a threshold of *∼* 4.7 (mol *·* min)*^−^*^1^, the *ZAP_active_* output jumps from 0 to a higher steady state of *∼*25%. This threshold is also seen with other parameters within the model, as shown with *k*4*f* (gray).

**Figure 2:**
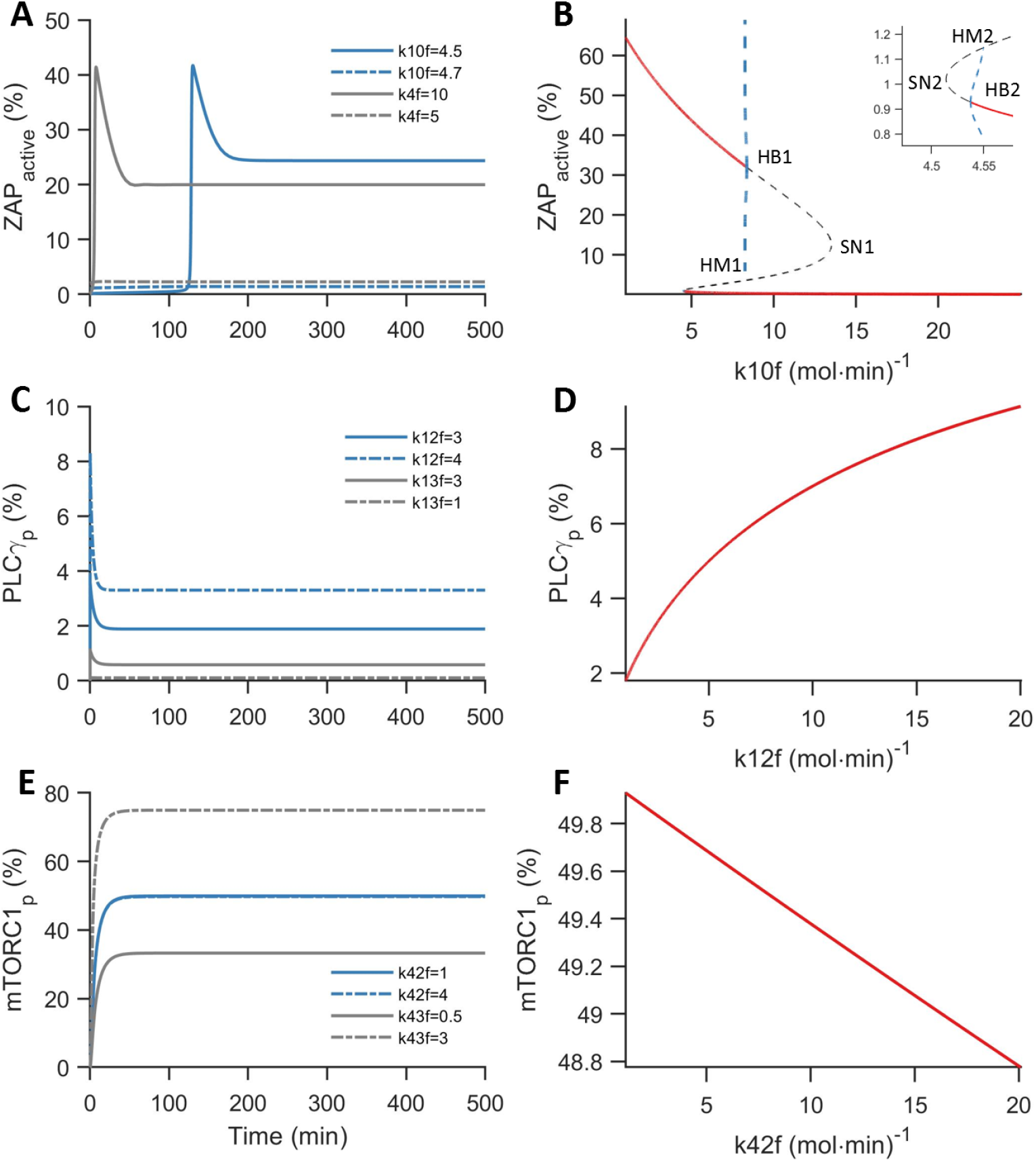
(A) Time series simulations of *ZAP_active_*, the output of the ZAP axis, at different values of *k*10*f* and *k*4*f* specified in the legend, where *k*10*f* and *k*4*f* are the parameters for *SHP* 1 phosphorylation, *SFK^∗^* and *ZAPp* binding, respectively. (B) One-parameter bifurcation of *ZAP_active_* with respect to *k*10*f*. Red (gray) solid (dashed) lines represent stable (unstable) branches of equilibria, while blue dashed lines represent envelopes of unstable limit cycles emerging from the Hopf bifurcations: HB1 and HB2, and terminating at the homoclinic bifurcations: HM1 and HM2. SN1 and SN2 are saddle-node bifurcations. (C) Time series simulations of *PLCγp* as a percentage relative to its total concentration: *PLCγ* + *PLCγp*, at different values of *k*12*f* and *k*13*f* specified in the legend, where *k*12*f* and *k*13*f* are the forward rate constants of *LAT* and *PLCγ* phosphorylation, respectively. (D) One-parameter bifurcation of *PLCγp* percentage with respect to *k*12*f*. Red line represents branch of stable equilibria. (E) Time series simulations of *mTORC*1*^∗^* as a percentage relative to its total concentration: *mTORC*1 + *mTORC*1*^∗^*, at different values of *k*42*f* and *k*43*f* specified in the legend, where *k*42*f* and *k*43*f* are the forward rate constants of *RhebGDP* and *mTORC*1 activation, respectively. (D) One-parameter bifurcation of *mTORC*1*^∗^* percentage with respect to *k*42*f*. Red line represents branch of stable equilibria. All time series simulations and bifurcation diagrams are plotted with *TCRp* = 27000 mol; for all other parameter values, see Table 6.1.

To further explore the dynamics of the model in more detail, a one-parameter bifurcation analysis of *ZAP_active_*with respect to *k*10*f* is conducted, as shown in Fig. 2(B). Branches of stable (unstable) steady states of the model are shown in red solid (gray dashed) lines. The gray dashed line exhibits loop points that represent saddle node bifurcations (SN1 and SN2). The stable and unstable branches are linked by Hopf bifurcations (HB1 and HB2) where envelopes of unstable limit cycles (blue dashed lines) emerge and terminate at homoclinic bifurcations (HM1 and HM2, respectively). As *k*10*f* decreases, a jump in the *ZAP_active_*steady state occurs upon crossing the HB2 bifurcation point. We also note that there is a bistability region within which the two branches of stable steady states overlap (shown by the region where the two red solid lines cover the same range of *TCRp* parameter values). This region is bounded by HB1 and HB2. Since the Hopf bifurcations produce envelopes of unstable limit cycles, periodic solutions are not observed when transitioning from one stable branch of steady states to the other. Additionally, since the bistable regime is bounded by the two Hopf bifurcations HB1 and HB2, rather than the two saddle nodes SN1 and SN2, as typically seen in traditional bistable switches, the bistable regime and the hysteresis of the model are quite limited in range. This bifurcation diagram suggests that certain molecular players within the ZAP axis are quite important in modulating its response to a vaccine input; however, it remains unclear which specific parameters within the axis are most sensitive to perturbations in their values.

Extending this analysis to the LAT and CD28 axes indicates that their dynamics are not as complex. Indeed, parameter sweeps and bifurcation analyses reveal a monostable behaviour across all the parameters within these two subsystems, suggesting that the LAT and CD28 axes do not exert the same level of influence on the overall behaviour of the system as the ZAP axis. This is demonstrated in Fig. 2(C,E), where the time series simulations of *PLCγp* and *mTORC*1*^∗^* percentages (relative to their individual total concentrations) from the LAT and CD28 axes, respectively, are plotted at different parameter values of *k*12*f* and *k*13*f* (denoting the forward rates of *LAT* and *PLCγ* phosphorylation, respectively) and at different parameter values of *k*42*f* and *k*43*f* (denoting the forward rates of *RhebGDP* and *mTORC*1 activation, respectively) provided in the legends; in these two cases, no threshold-dependent sudden jumps in their steady state output dynamics are observed. This is further confirmed in Fig. 2(D,F) showing a monotonically varying branch of stable equilibria in the bifurcation diagrams of *PLCγp* and *mTORC*1*^∗^* percentages with respect to *k*12*f* and *k*42*f* , respectively. This emphasizes the limited significance of these parameters in defining the dynamics of the LAT and CD28 axes output variables.

These results thus suggest that a circadian rhythm acting on signaling molecules within the ZAP axis would have the largest impact on model dynamics. This is validated in the next section when performing sensitivity analysis on the parameters of the ZAP axis.

##### 3.2.2. Parameters of ZAP 70 and SHP 1 activation are the most sensitive to perturbation

To identify sensitive signaling molecules within the ZAP axis, partial rank correlation coefficient (PRCC) of each parameter within a given axis is calculated (Marino et al., 2008). A partial correlation characterizes the relationship between a parameter *p_i_* and the output variable, while discounting the effects of other parameters in the model. This is repeated for every activation parameter *p_i_* within the ZAP axis, and is done using a MATLAB package written by Marino et al. (Marino et al., 2008), which incorporates Latin Hypercube Sampling (LHS) to generate parameter sets whose partial correlation coefficients are calculated. The results of this are shown in Fig. 3. From this plot, we designate *k*4*f, k*5*f, k*8*f* 2*, k*9*f* 1 and *k*10*f* as sensitive parameters, i.e. with correlation coefficients *> |*0.6*|*. This threshold correlation coefficient represents 80% of the correlation coefficient of *k*10*f* (whose correlation coeficient is 0.72), where *k*10*f* denotes the rate of activation of the *SHP* 1 molecule. Since *SHP* 1 has been suggested to possess a circadian component in its regulation (Nobis et al., 2019b), we assume that the other circadian-affected parameters of the model should posses a level of sensitivity equivalent or close to that possessed by *SHP* 1 activation through *k*10*f*. Interestingly, those parameters identified with this approach are involved in the formation of *ZAP_active_*, with *k*4*f* and *k*5*f* denoting the forward and backward rates of *SFK^∗^*-*ZAPp* phosphorylation, and *k*8*f* 2 and *k*9*f* 1 denoting the formation rates of *ZAP ^∗^* and *ZAPp*, respectively. Additionally, this sensitivity analysis demonstrates the importance of negative feedback in affecting *ZAP_active_* through *SHP* 1*^∗^*. Since *k*10*f* is the primary negative feed-back parameter implicated by this analysis, it will be our target parameter in the single-parameter analysis conducted in the following section.

**Figure 3:**
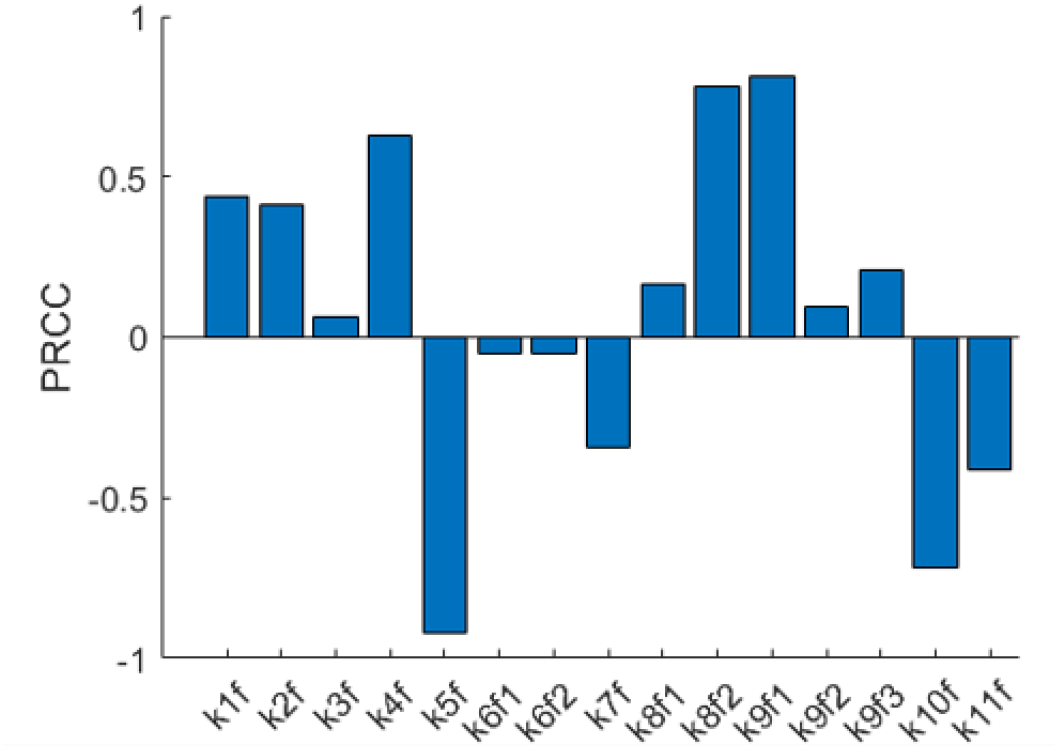
Partial rank correlation coefficients (PRCC) of the activation parameters of the ZAP axis when correlated to *ZAP_active_* output. A higher PRCC value indicates a higher correlation between values of that parameter with *ZAP_active_*(positively correlated if PRCC *>* 0 and negatively correlated if PRCC *<* 0). The following parameters *k*4*f, k*5*f, k*8*f* 2*, k*9*f* 1 and *k*10*f* are identified as sensitive with correlation coefficients *> |*0.6*|*.

#### 3.3. The impact of vaccination on the steady state dynamics of the model

Implementing the periodic forcing function defined by Eqn. 1 on k10f (the rate of activation of the SHP 1 molecule) will allow this parameter to exhibit a 24 hr cycle, mimicking the influence of a circadian rhythm. This is shown in Fig. 4(A), where *k*10*f_t_* is plotted according to the circadian time of day, with [0, 12] hr representing day-time and [12, 24] hr representing night-time. The amplitude of this diurnal variation in *k*10*f_t_* varies according to *α_k_*_10_*_f_* denoting the coefficient of variation; when *α_k_*_10_*_f_* = 0.5 (solid line), larger oscillations are produced compared to *α_k_*_10_*_f_* = 0.25 (dashed line). The respective day and night time maxima and minima of *k*10*f_t_* are identified by an orange (yellow) and blue (light blue) dot for each value of *α_k_*_10_*_f_* , respectively. Recall that (see Fig. 2) the bifurcation diagram of the model exhibits a bistable switch with two saddle-nodes (SN1 and SN2) and two Hopf bifurcations (HB1 and HB2) at which two envelopes of unstable periodic orbits emerge and eventually terminate at homoclinic bifurcations (HM1 and HM2, respectively). Superimposing the orange and blue dots representing the maxima and minima of *k*10*f_t_* when *α_k_*_10_*_f_* = 0.5 on this bistable switch, as shown in Fig. 4(B), the orange dot ends up lying on the upper branch of stable equilibria (to the left of HB1), while the blue one ends up lying on the lower branch (to the right of HB2). This means that during a 24 hr day-night cycle in *k*10*f_t_*, the solution trajectory of *ZAP_active_* will oscillate between these two dots that represent the extrema of *k*10*f_t_* and cause it to jump back and forth between the two stable branches of equilibria. The jumping to the lower branch will occur during the transition from daytime to nighttime when the trajectory moves rightward along the upper branch of *ZAP_active_* stable equilibria until it crosses HB1, leading the trajectory to jump to the lower stable branch and causing *ZAP_active_* to decrease. Conversely, during the transition from nighttime to daytime, the trajectory will move leftward along the lower branch of *ZAP_active_*stable equilibria until it crosses HB2, leading the trajectory to jump to the upper stable branch and causing *ZAP_active_*to increase. Thus, a circadian rhythm on ZAP axis activation parameters can have a drastic effect on *ZAP_active_* output. This drastic change in dynamics occurs because of the presence of the bistable switch with its two Hopf bifurcations: HB1 & HB2, lying within the parameter space afforded by *α_k_*_10_*_f_* = 0.5. This cycling between the two branches of equilibria every 12 hr suggests that this circadian rhythm can cause periodic oscillations in *ZAP_active_* with large amplitudes. This then poses the question of how such mechanism interacts with vaccines?

**Figure 4:**
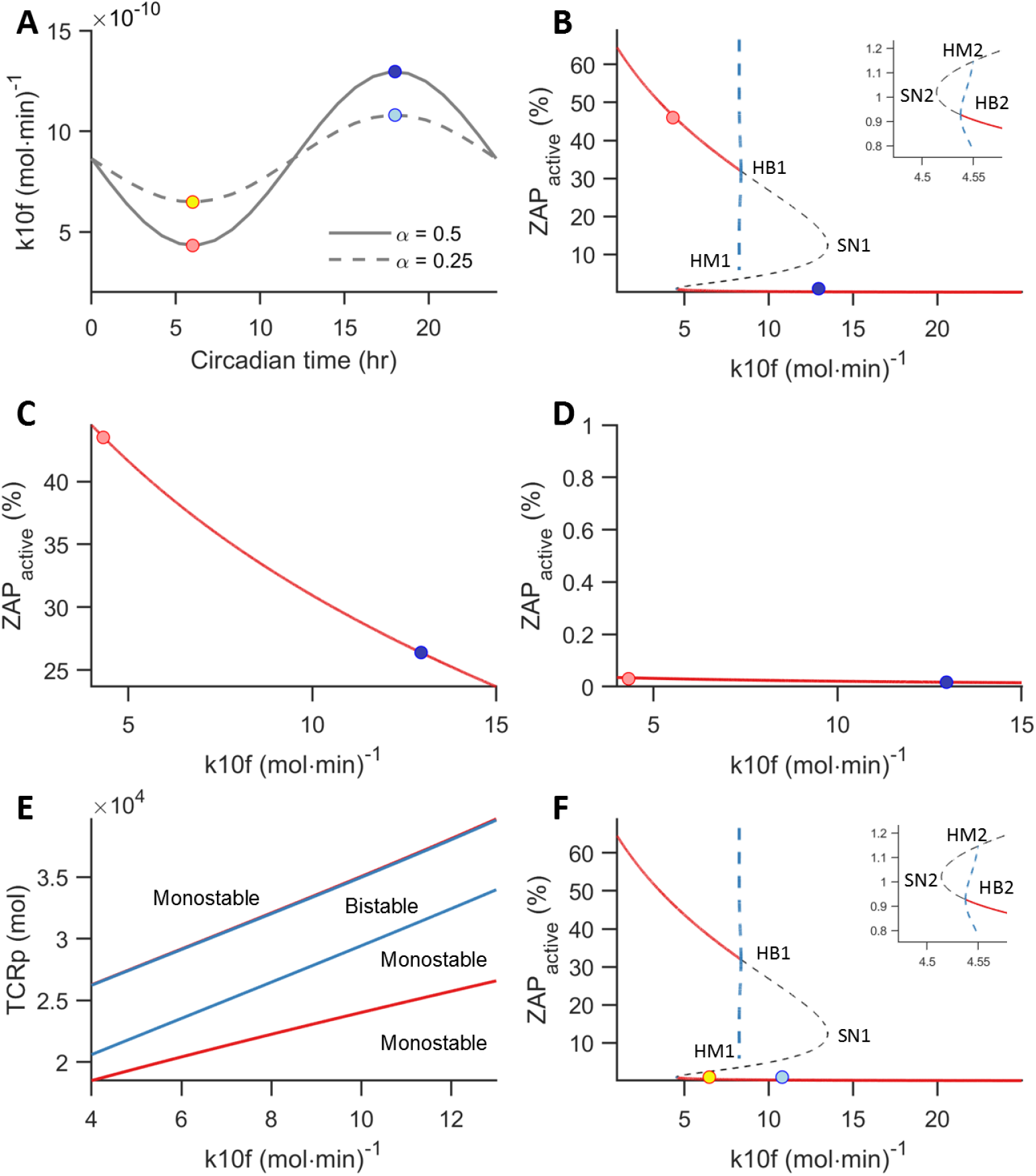
(A) Time series plots of *k*10*f_t_* when *α_k_*_10*f*_ = 0.5 (solid line) and *α_k_*_10*f*_ = 0.25 (dashed line) showing 24 hr cycles with different amplitudes. Orange (yellow) and blue (light blue) dots show the diurnal maxima and minima of *k*10*f_t_* when *α_k_*_10*f*_ = 0.5 (*α_k_*_10*f*_ = 0.25). (B) One-parameter bifurcation of *ZAP_active_* with respect to *k*10*f* at *TCRp* = 27000 mol. Details of the bifurcation diagram are described in Fig. 2. Orange and blue superimposed dots represent the respective day and night extrema of *k*10*f_t_* identified in (A). Note the abrupt change in steady state upon crossing HB1 and HB2. (C), (D) One-parameter bifurcation of *ZAP_active_* with respect to *k*10*f* at *TCRp* = 40000 mol (C) and *TCRp* = 10000 mol (D). Note the monostable response across the *k*10*f_t_* parameter space. (E) Two-parameter bifurcation of *ZAP_active_* with respect to *k*10*f* and *TCRp*. Red lines denote continuations of saddle-node bifurcations (SN1 and SN2), while blue lines denote continuations of the Hopf bifurcations (HB1 and HB2). Bistability is bounded by HB1 and HB2. (F) one-parameter bifurcation of *ZAP_active_* at *TCRp* = 27000 mol as in (B), but with *α_k_*_10*f*_ = 0.25; the day-night extrema identified in (A) are superimposed as yellow and light blue dots.

To answer this question, we will investigate how the impact of a circadian rhythm on CD8^+^ T cell activation is affected by the *TCRp* (the number of phosphorylated TCRs), which we assume to be an indicator of vaccine strength. Thus, an elevated *TCRp* is assumed to demonstrate a vaccine presentation to the CD8^+^ T cell, and a higher vaccine load is similarly assumed to trigger a higher number of phosphorylated TCRs, thus leading to a larger *TCRp* model input. This is first done by considering two other values of *TCRp*: one that is higher than the one used in Fig. 4(A), namely, *TCRp* = 40000 mol, and another that is lower, i.e., *TCRp* = 10000 mol, and then plotting the one-parameter bifurcations of *ZAP_active_* with respect to *k*10*f* at these two *TRCp* values as shown in Fig. 4(C,D), respectively. By superimposing the orange and blue dots representing the day and night extrema of *k*10*f_t_* when *α_k_*_10_*_f_* = 0.5 on these two bifurcation diagrams, we see that in both realizations, no Hopf or saddle node bifurcations end up separating the two dots to allow for drastic time-of-day dependent changes in *ZAP_active_* seen in Fig. 4(B). Indeed, when *TCRp* = 40000 mol, the two dots become part of the upper branch of stable equilibria, producing an elevated output of *ZAP_active_* independent of time of day, whereas when *TCRp* = 10000 mol, the inverse is seen, with the two dots lying on the lower branch of stable equilibria, producing baseline output of *ZAP_active_* independent of time of day. Thus, there seems to be an intermediate range of *TCRp* within which this circadian rhythm can produce the largest day-night variation in *ZAP_active_*. This relationship between *TCRp* and *k*10*f* is further demonstrated by plotting the two-parameter bifurcation of *ZAP_active_* with respect to both *TCRp* and *k*10*f* as shown in Fig. 4(E). The region bounded by the Hopf bifurcations (blue lines) demarcates the bistable regime, whereas the upper and lower regions denote the monostable elevated and baseline states of the model, respectively. Notice how the bistable regime shifts horizontally as the value of *TCRp* is varied, indicating that there is an intermediate range of *TCRp* within which the largest day-night variation in *ZAP_active_* can be generated. This means that the level of TCR phosphorylation, or level of vaccine load, can alter *TCRp* and cause the regime of bistability to shift accordingly.

Now that we have explored how differing vaccine loads may affect bistability with respect to *TCRp* and *k*10*f* , we now examine how changing the strength of this circadian rhythm through *α_k_*_10_*_f_* can affect outcomes. By tak-ing *TRCp* = 27000 mol and setting *α_k_*_10_*_f_* = 0.25 to produce the two extrema of *k*10*f_t_*identified by the yellow and light blue dots in Fig. 4(A), we find that superimposing these dots on the bifurcation diagram of *ZAP_active_* with respect to *α_k_*_10_*_f_* (see Fig. 4(F)) places them on the lower stable branch of equilibria. According to this model realization, solution trajectories are unable to cross the Hopf bifurcation HB2 and solely oscillate along this lower branch with very low amplitude, i.e., without exhibiting drastic day-night shift in *ZAP_active_*. Such a realization thus indicates that if the influence of this circadian rhythm is not strong enough, then an intermediate level of vaccination may still not be able to generate large amplitude day-night cycle in *ZAP_active_*.

Taken together, these results demonstrate that the day-night variation in signaling is dependent on both having a sufficiently large *TCRp* input, as well as on possessing a sufficiently high *α_k_*_10_*_f_* , such that the *k*10*f_t_* is able to cross HB1 and HB2 of the model. This further highlights the complex relationship between vaccine load and the strength of circadian modulation when mediating CD8^+^ T cell response upon vaccination. Further experimental work is needed to validate the extent of this relationship.

#### 3.4. Impact of circadian coupling on temporal evolution of CD8^+^ T cell signaling

Now that we have determined the underlying steady state dynamics of circadian modulation of CD8^+^ T cell activation with different vaccine loads, we explore how the time series of the ZAP axis is affected by *k*10*f_t_* and the potential physiological implications of this on CD8^+^ T cell signaling. To do so in the context of vaccination, we define two possible states that can be attained by the model: *S*_0_ and *S*_1_. *S*_0_ represents the starting state of the model (pre-vaccination) with an initial amount of *TCRp* given by *TCRp*_0_ = 100 mol; a nonzero basal level of *TRCp*_0_ is chosen for this state *S*_0_ to account for pMHC-TCR contacts with self or non-cognate antigens. *S*_1_, on the other hand, denotes the post-vaccination state and corresponds to when *TCRp >>* 10000 mol (*>> TCRp*_0_), i.e., when the CD8^+^ T cell has been presented with a vaccine. The effect of a circadian rhythm on the time series of the ZAP axis in each of these states is then analyzed (see Fig. 5) by exploring the effects of a *k*10*f* -induced circadian rhythm on both *S*_0_ or *S*_1_.

**Figure 5:**
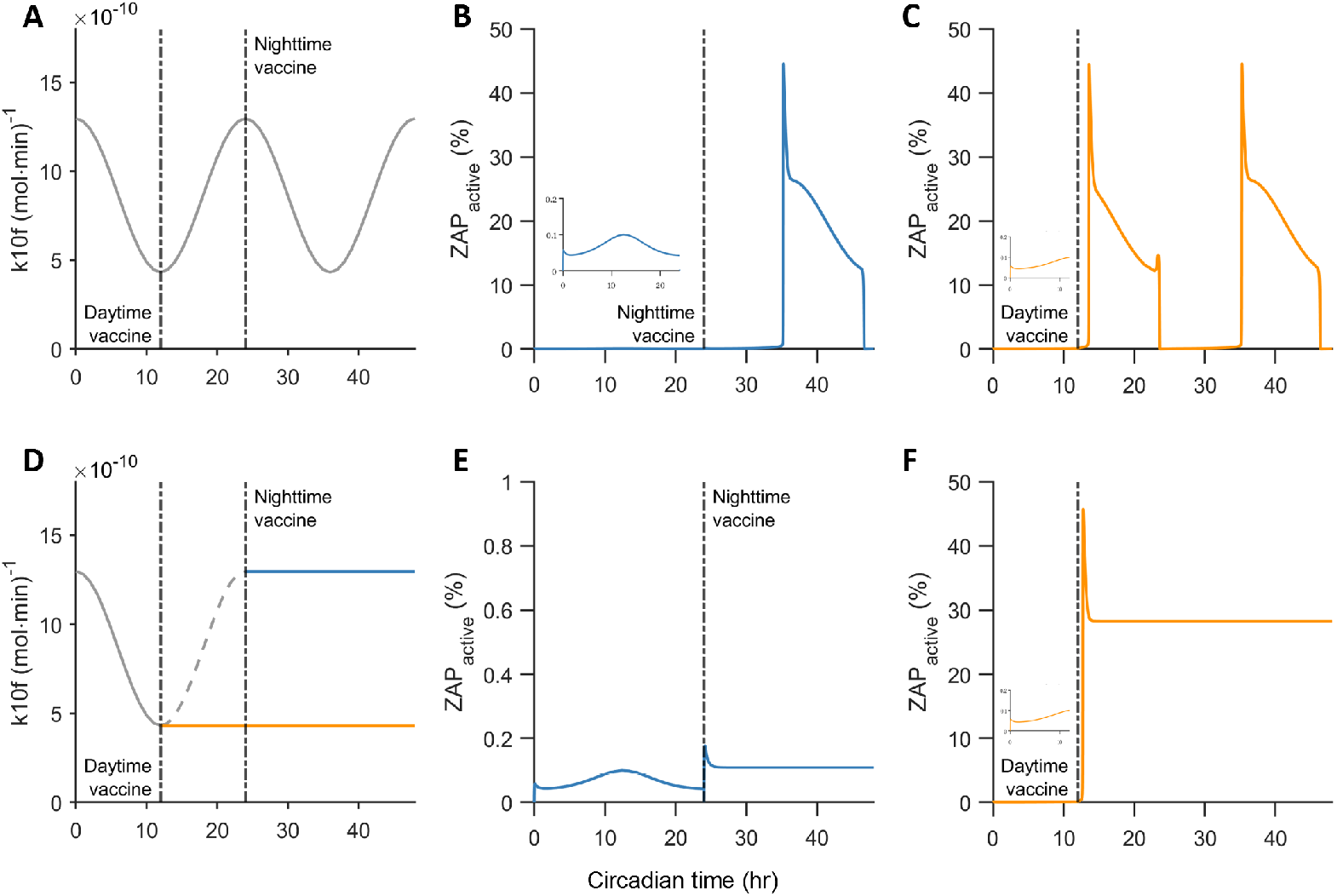
(A) Time series plot of *k*10*f_t_* when *α_k_*_10*f*_ = 0.5 showing 24 hr cycles. Dashed-dotted lines indicate the moments of vaccine applications at daytime (*t* = 12 hr) and at nighttime (*t* = 24 hr). (B) Time series simulations of *ZAP_active_* pre and post vaccination applied at nighttime (*t* = 24 hr). Inset: magnification of the time series simulations prio to vaccination to highlight the small amplitude oscillations associated with the state *S*_0_. (C) Time series simulations of *ZAP_active_*pre and post vaccination applied at daytime (*t* = 12 hr). Notice the repetitive large amplitude oscillations associated with the state *S*_1_ produced post vaccination at nighttime and daytime; the onset of the large amplitude oscillations in the former case (nighttime vaccination) is delayed relative to the latter case (daytime vaccination). (D) Time series plot of *k*10*f_t_* when *α_k_*_10*f*_ = 0.5 showing 24 hr cycles that get interrupted upon vaccination at daytime (orange; 12 hr) and nighttime (blue; 24 hr). When vaccination is applied in each case, *k*10*f_t_* is then kept steady at its respective value. (E) Time series simulations of *ZAP_active_* pre and post vaccination applied at nighttime (*t* = 24 hr). (F) Time series simulations of *ZAP_active_*pre and post vaccination applied at daytime (*t* = 12 hr). Inset: magnification of the time series simulations prior to vaccination to highlight the small amplitude oscillations associated with the state *S*_0_. Notice the much larger response associated with *S*_1_ produced post vaccination at daytime compared to nighttime, as well as the absence of oscillations post vaccination in both cases.

Imposing a 24-hr circadian cycle on *k*10*f* in the form of a sine wave with *α_k_*_10_*_f_* = 0.5 as in Fig. 5(A) and initiating model simulations from *S*_0_ induce a repetitive very small-amplitude oscillations is *ZAP_active_* (Fig. 5(B, inset)). Although *TRCp* = 10000 mol in Fig. 5(D), the monostable dynamics detected in this figure is responsible for generating these small amplitude oscillations that shuttle *ZAP_active_* between the orange and blue dots. This is because the bifurcation diagram at *S*_0_ is essentially the same as the one in Fig. 5(D). However, when TCRs become phosphorylated, due to vaccine application, the state of the system switches to *S*_1_. By setting *TCRp* = 29000 mol in *S*_1_, *ZAP_active_* then produces repetitive large amplitude oscillations regardless of the time of vaccination (Fig. 5(B,C)), with the daytime vaccination (at *t* = 12 hr) producing a faster response compared to the nighttime vaccination (at *t* = 24 hr). These large amplitude oscillations are reminiscent to those cycles between the orange and blue dots highlighted previously in Fig. 5(B); solution trajectories, in this case, cross the two Hopf bifurcations back and forth and jump between the branches of stable equilibria. Given that such oscillations indicate repetitive activation and deactivation of T cells, then this would preclude them from being physiological.

To address this latter issue, we hypothesize that upon vaccine application the effect of the circadian oscillations on *k*10*f* is significantly attenuated with a much lower value for *α_k_*_10_*_f_* . To test this hypothesis, we set *α_k_*_10_*_f_* = 0 right when a vaccine is applied at both daytime (at *t* = 12 hr) and nighttime (at *t* = 24 hr) as shown in Fig. 5(D). This is done by freezing (i.e., fixing) the value of *k*10*f* at its respective value upon vaccine onset. Even though vaccine application still transitions the system from the state *S*_0_ to the state *S*_1_, repetitive large amplitude oscillations no longer occur only small amplitude oscillations in *ZAP_active_* is seen prior to vaccination due to the circadian clock as described before for *S*_0_. Interestingly, nighttime vaccination causes a very moderate increase in *ZAP_active_* (Fig. 5(E)), whereas daytime vaccination causes a significant increase in *ZAP_active_* (Fig. 5(F)), reaching a significantly elevated steady state due to the transition to the upper branch of stable equilibria highlighted in Fig. 5(B). The lack of oscillations in *k*10*f_t_* post vaccination prevents the solution trajectory from crossing HB1 and returning back to the lower branch. These results thus corroborate the experimental finding that daytime vaccination is more effective in triggering T cell activation, and suggests that for this to be true, the diurnal effect of a circadian rhythm on early events of CD8^+^ T cell signaling is significantly attenuated post vaccination. Based on these observations, we conclude that this circadian rhythm primes or facilitates the activation of näıve CD8^+^ T cells to produce the day-night variation in *ZAP_active_* output, rather than directly controlling it.

#### 3.5. Characterizing the filtering properties of the ZAP axis

Throughout this study, we have explored the implications of a circadian rhythm on the ZAP axis, which we have found to be the primary driver of model dynamics with its intrinsic bistable switch. Here, we expand this study by exploring how the effects of a circadian rhythm acting directly on the ZAP axis is transmitted to the downstream LAT and CD28 axes within the model in the absence of a vaccine input, i.e., in the *S*_0_ state. Doing so would indicate whether a circadian rhythm within the ZAP axis is capable of producing a day-night variation in the signaling of other downstream molecular players associated with these two axes. If not, this may suggest that those molecular players may be independently regulated by the circadian clock.

To explore how the influence of the circadian oscillation within the ZAP axis in the absence of vaccination is transmitted to other axes of the model, we now consider the expanded model of Fig. 1 that includes the LAT and CD28 axes. In order to quantify how the circadian-dependent oscillations of *k*10*f_t_* are transmitted to other variables in these two axes, we will use the ’oscillatory contribution’ ratio, *ω_i_*, for every variable (molecular player) *i* of the model. This *ω_i_* is defined as the relative effect of the oscillation in *k*10*f_t_* to the mean of the variable *i* at steady state. It is given by

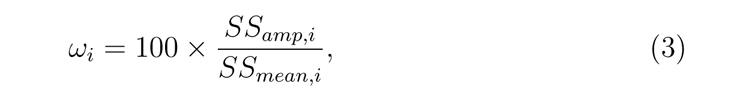

where *i* represents the variable of interest, *SS_amp,i_* represents the amplitude of the steady state oscillations, and *SS_mean,i_*represents the mean of the variable. A larger *ω_i_* indicates that a larger proportion of a variable’s steady state can be explained by *k*10*f_t_*, whereas a smaller *ω_i_* indicates that the circadian-dependent oscillations in *k*10*f_t_* minimally alter the steady state of the variable *i*. To quantify *ω_i_*, we will always assume that *k*10*f_t_* is oscillatory with a circadian coefficient of *α_k_*_10_*_f_* = 0.5. Since the model is in *S*_0_, this value of *α_k_*_10_*_f_* is chosen as it creates the largest variation in steady state amplitude, allowing us to more easily visualize how a circadian rhythm is filtered through the model, without the possibility of jumping to an upper, ’activated’, steady state branch.

By plotting the magnitude of *ω_i_*for the major variables within the ZAP axis, as shown in Fig. 6 (blue bar), we find that *SHP* 1*_active_* is associated with the largest *ω_SHP_* _1_, while the other variables (including *i* = *SFK^∗^, CSK^∗^, CBPp*) exhibit a reduced *ω_i_*. This is expected since a periodic forcing on *k*10*f* directly regulates SHP1 activation, altering *SHP* 1*^∗^*. For the variables within the LAT axis, on the other hand, Fig. 6(B) shows that the magnitude of *ω_i_* (*i* = *LATp, PLCγp, SOSb*) are lower than those for the key molecular players within the ZAP axis: *ZAP_active_* and *SHP* 1*^∗^*, suggesting that some of the *k*10*f* -induced oscillation is filtered out between axes. However, the oscillations remain identifiable within the axis, indicating that a circadian rhythm in the ZAP axis is sufficiently transmitted to the LAT axis. Interestingly, in the CD28 axis, Fig. 6(C) shows that while the oscillation from *k*10*f_t_* is adequately transmitted to many of the signaling molecules (including *i* = *PI*3*K^∗^, PDK*1*^∗^, AKT ^∗^, mTORC*2*^∗^*) with moderate values of *ω_i_*, it is completely filtered out for *i* = *TSC*1*−* 2*, RhebGTB, mTORC*1*^∗^* (i.e., *ω_i_* = 0 for all of these cases). This suggests that oscillations in *k*10*f_t_* directly influencing the *ZAP* axis may not be able to produce the experimentally observed oscillations within *mTORC*1*^∗^* and its immediate precursors (Nobis et al., 2019b), despite producing them in the other signaling molecules of this axis.

**Figure 6:**
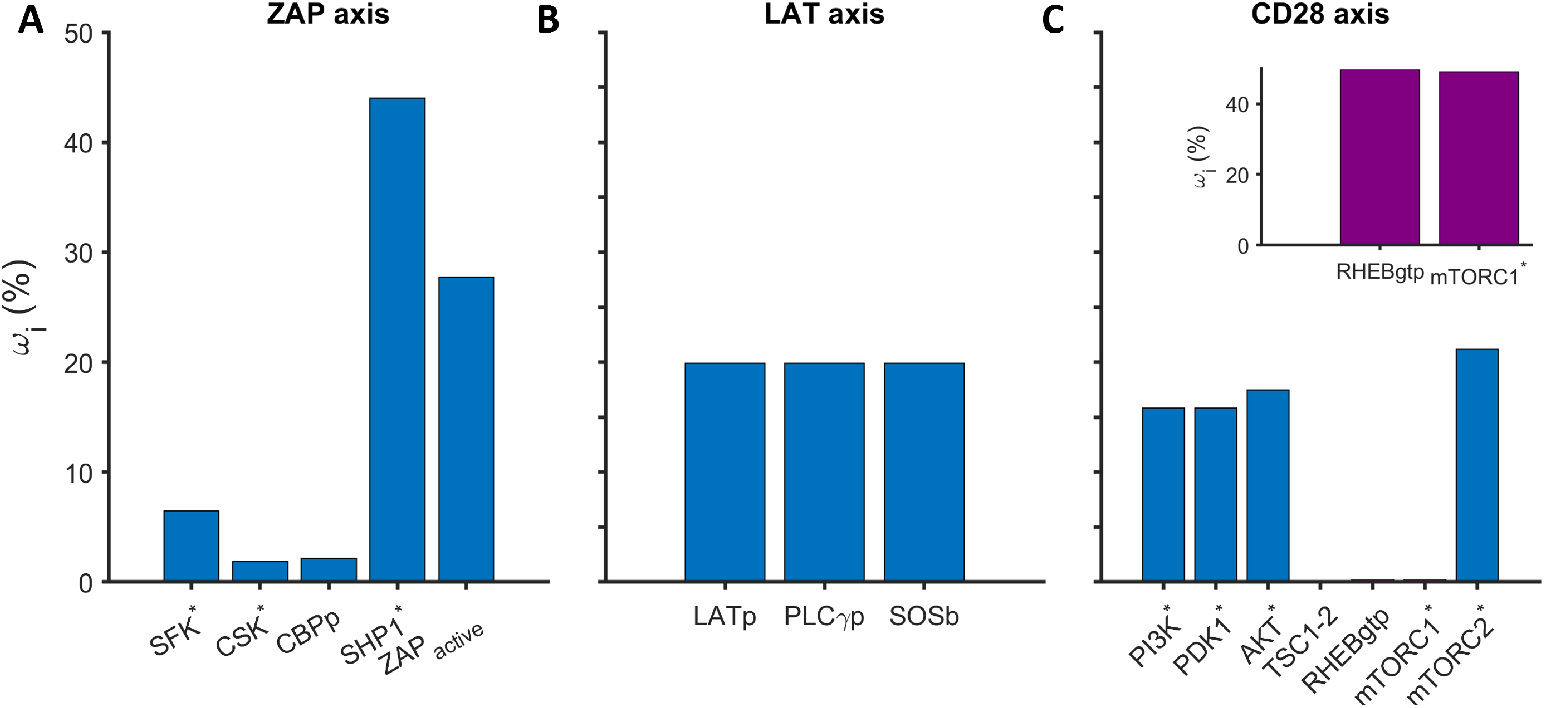
Bar plots of the magnitude of the oscillatory contribution *ω_i_* for different molecular players within the (A) ZAP, (B) LAT, and (C) CD28 axes when *k*10*f_t_* is oscillatory with *α_k_*_10*f*_ = 0.5. The larger the *ω_i_* value, the stronger the influence of the circadianinduced oscillations have on the steady state level of the molecular player *i* of interest. Inset in panel (C): Bar plots of the magnitude of the oscillatory contribution *ω_i_* for the molecular players *i* = *RhebGTP* and *mTORC*1*^∗^* within the CD28 axis when *k*42*f_t_* is oscillatory with *α_k_*_42*f*_ = 0.5.

This was resolved by imposing a periodic forcing component, representing the influence of a circadian rhythm, directly on *RhebGTP* activation parameter *k*42*f* with *α_k_*_42_*_f_* = 0.5; this resulted in a high *ω_RhebGT P_* and *ω_mT ORC_*_1_*_active_* of 49% and 49.5%, respectively, as shown in Fig. 6(C, inset). Curiously, adding a periodic forcing onto any of the more upstream signaling molecules within the CD28 axis, such as AKT, PI3K or even TSC1-2, was not able to produce oscillations in *RhebGTP* and *mTORC*1*^∗^*, further indicating the presence of an independent circadian regulation of *RhebGTP* and *mTORC*1*^∗^*. Since *S*6 (a molecular derivative of *mTORC*1), has been experimentally shown to have a circadian-influenced day-night variation (Nobis et al., 2019b), these modeling results suggest that a circadian rhythm within the ZAP axis may not be sufficient to explain these downstream oscillations, pointing to *mTORC*1 and its related players being secondary, independent targets of the molecular circadian clock.

## 4. Population model approach

### 4.1. Methodology

To better understand CD8^+^ T cell responses to circadian oscillations and vaccines, it is imperative to obtain a broader view of model dynamics at the T cell population level. We do so by simulating the steady state dynamics of *n* = 1000 model realizations of the ZAP axis, each with a randomly selected parameterization of the model. Similarly to the single-cell approach, the oscillatory regime of the model was excluded using the framework described in Appendix 6.3. This has allowed us to confine parameter sampling of each realization to the non-oscillatory regime only.

### 4.2. One-parameter macro-bifurcation analysis

We begin our population level analysis by first targeting one specific parameter, namely, *k*10*f_k_*. This is done by randomly assigning values for this parameter within the range [0.5 *× k*10*f*_0_, 1.5 *× k*10*f*_0_] for each model realization *k*, where *k*10*f*_0_ denotes the default parameter value of *k*10*f* listed in Table 6.1. Doing so generates heterogeneity between each model realization through this parameter. These model realizations are then all integrated from the same initial conditions at increasing values of *TRCp ∈* [1, 3.5] *×* 10^4^ mol and the steady state values of *ZAP_active_* are plotted with respect to *TRCp* (along the x-axis) as shown in Fig. 7(A). Each point on this graph represents a steady state value of *ZAP_active_* at a given *k*10*f_k_* within the previously defined range. Doing so provides a snapshot of 1000 steady states of the model, each at a different, randomly selected, circadian time defined by the value of *k*10*f_k_ ∈* [0.5 *× k*10*f*_0_, 1.5 *× k*10*f*_0_] (recall that we assumed that this circadian rhythm affects the value of *k*10*f* when we defined *k*10*f_t_*). Plotting the steady states of these 1000 realizations with respect to *TCRp*, the surrogate for vaccine strength, highlights how the model output varies under differing vaccine loads. This produces what looks like a one-parameter bifurcation diagram that encompasses not only the effects of this circadian rhythm on T cell activation through *k*10*f* , but also the effects of vaccine strength through *TRCp*. We refer to this plot as a ’macro-bifurcation’ diagram.

**Figure 7:**
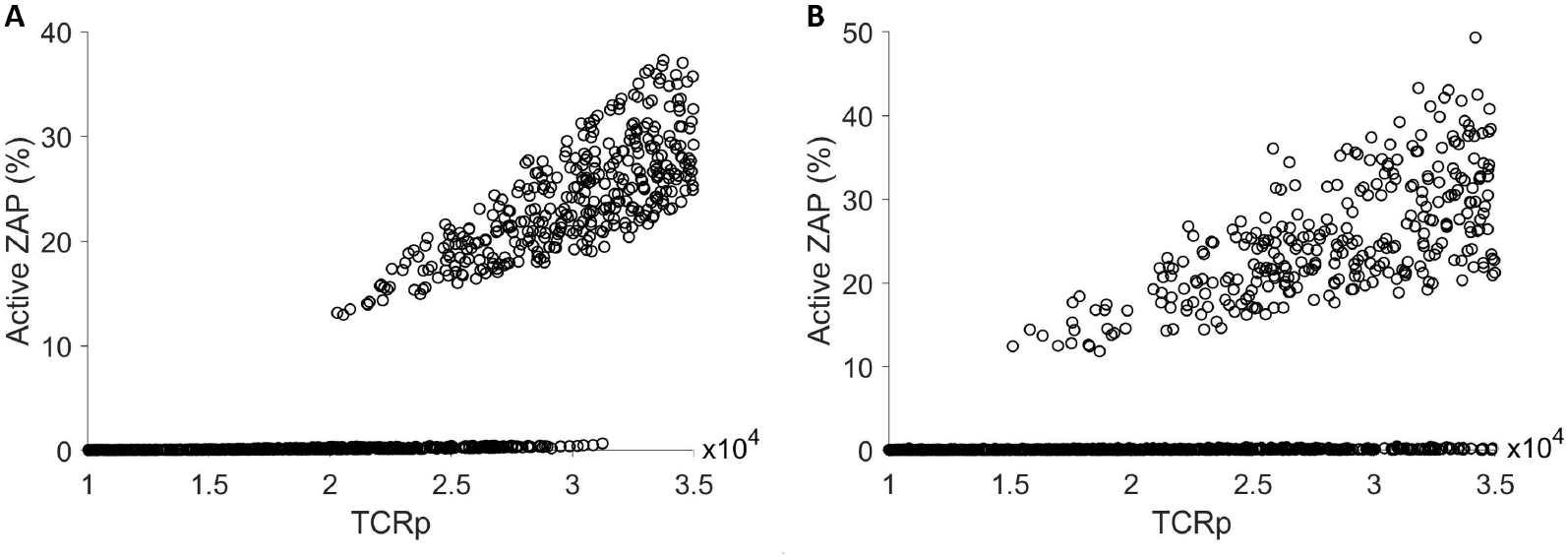
Macro-bifurcation diagram of *ZAP_active_* with respect to *TRCp* when various combinations of circadian-affected parameters are assigned random values within their respective ranges: (A) *k*10*f* is randomly assigned values within the range [0.5 *× k*10*f*_0_, 1.5 *× k*10*f*_0_]. (B) *k*4*f, k*9*f* 1*, k*8*f* 1*, k*11*f, k*7*f, k*10*f* are randomly assigned values within the ranges [0.5 *× par*_0_, 1.5 *× par*_0_], where *par*_0_ = *k*4*f*_0_*, k*9*f* 1_0_*, k*7*f*_0_*, k*10*f*_0_*, k*11*f*_0_*, k*8*f* 1_0_.

According to this macro-bifurcation diagram, we can split the parameter space of the model into 3 regions labeled insensitive, sensitive and elevated. This distinction depends on the distribution of *ZAP_active_* at each value *TCRp*. The insensitive region is the range of *TCRp ∈* [1, 2]*·*10^4^ mol, where *ZAP_active_*is purely in its inactive state, regardless of the parameterization of *k*10*f*. The sensitive region, on the other hand, corresponds to the range of *TCRp ∈* [2*, ∼* 3.10] *·* 10^4^ mol, where *ZAP_active_* can be either elevated or not, depending on the value of the parameter *k*10*f_t_*. We denote this region as possessing ’macro-bistability’, as the magnitude of its output depends on the value of the circadian-affected parameter *k*10*f*. Finally, the elevated region is the one that corresponds to the range *TCRp ∈* [*∼* 3.10, 3.5] *·* 10^4^ mol, where *ZAP_active_* is purely elevated, regardless of the value of *k*10*f_t_*.

### 4.3. Multi-parameter macro-bifurcation analysis

Unique to the macro-bifurcation analysis of the previous section is the ability to expand it to examining *n* parameters simultaneously, an advantage not afforded by the standard bifurcation analysis that is typically limited to at most 2 parameters (as in Fig. 4E). To harness this advantage, we now include variability not only in *k*10*f* , but also in the positive feedback parameters *k*4*f, k*9*f* 1, negative feedback parameters *k*7*f, k*11*f* and our manually derived threshold parameter *k*8*f* 1 (see Appendix 6.3.2). We have chosen these parameters because they can act as possible molecular targets of a circadian rhythm in early CD8^+^ T cell signaling and because they were a subset of ’sensitive’ parameters identified by our sensitivity analysis of Fig. 3 (for the full definition of *k*4*f, k*9*f* 1*, k*7*f, k*10*f, k*11*f* and *k*8*f* 1, please see Tables 6.1). Combining their effects in a macro-bifurcation diagram can thus provide a more physiological understanding of how a circadian rhythm can impact CD8^+^ T cell activation. Simulating 1000 realizations of the model, by randomly, and independently, sampling these parameters from their respective ranges defined by [0.5 *× par*_0_, 1.5 *× par*_0_], where *par* = *k*4*f, k*9*f* 1*, k*7*f, k*10*f, k*11*f, k*8*f* 1 and *par*_0_ is the default parameter value of each one of them (see Tables 6.1), and plotting the steady state values of *ZAP_active_*produced by the model with respect to *TCRp* produces the macro-bifurcation diagram shown in Fig. 7(B). Although the distinction between insensitive, sensitive and elevated regions remains regardless of the number of parameters included, the macro-bistable region of the model (the sensitive region) becomes more expanded (i.e., more sensitive) compared to that shown in Fig. 7(A): *TCRp ∈* [2, 3] *·* 10^4^ mol in panel (A) vs *TCRp ∈* [1.5, 3.5] *·* 10^4^ in panel (B). This demonstrates that our theoretical framework is maintained when adding additional parameters, and suggests that incorporating variability in additional parameters in the macro-bifurcation analysis increases the range of the macro-bistability by introducing more heterogeneity in CD8^+^ T cells. This increased heterogeneity due to the circadian rhythm may improve the sensitivity of CD8^+^ T cells at the population level, allowing them to activate at a lower vaccine load. Further analysis, coupled with experimental work, may help determine the mechanism by which these additional rate parameters interact with the circadian rhythm directly to produce this increased sensitivity, providing a more complete understanding of the complex population-level immune-circadian interaction.

## 5. Discussion

In this study, we provided a detailed analysis of the dynamics of the signaling molecules involved in producing the day-night variation seen in CD8^+^ T cells signaling. Additionally, we explored how this day-night variation in CD8+ T cells downstream of the T cell receptor (TCR) may be altered by different levels of TCR phosphorylation. This was done through a mathematical modeling approach, which provided flexibility in studying how changes in rate activation parameters can affect CD8^+^ T cell activation.

More specifically, a reference model for CD4^+^ T cell activation obtained from Zheng et al. (2005); Perley et al. (2014) was adapted to CD8^+^ T cell activation and used to conduct our analysis. This model was then split into 3 subsystems, and a dynamical systems analysis was conducted on each one to identify the main drivers of model dynamics. Through this analysis, the subsystem associated with *ZAP* 70 activation (ZAP axis) was shown to drastically affect the dynamics of the other two downstream LAT and CD28 axes by uniquely possessing a bistable switch. This is in agreement with prior modeling literature suggesting a bistability within ZAP70 signaling (Lipniacki et al., 2008). To further pinpoint potential molecular targets of a circadian rhythm within the ZAP axis, a global sensitivity analysis was conducted using Latin Hypercube Sampling (LHS) and Partial Rank Correlation Coefficient (PRCC) (Marino et al., 2008). This was done to identify how parameter perturbations within the ZAP axis can affect the dynamics of its output signal, namely *ZAP_active_*. Since this circadian rhythm was implemented in a parameter-specific manner, identifying the most sensitive parameters of the model provided insights into potential targets of the circadian clock in CD8^+^ T cell signaling. Through this analysis, we identified that *ZAP* 70 and *SHP* 1 activation parameters, namely *k*4*f, k*5*f, k*10*f, k*9*f* 1 and *k*8*f* 2 were the most sensitive to perturbations, conclusions that are aligned with the results of transcriptomic analyses of CD8^+^ T cells in mice (Nobis et al., 2019b).

To analyze the effects of this parameter-specific circadian coupling on model dynamics, a single-parameter bifurcation analysis approach was applied. Due to the bistable switch within the ZAP axis, we showed that, for a given *TCRp*, an elevated *ZAP_active_* was dependent on the value of the circadian-coupled bifurcation parameter *k*10*f_t_*, especially relative to its Hopf bifurcation points, thus producing a day-night variation. This day-night variation was also shown to be affected by the quantity of TCR phosphorylation, which we designate as the vaccine load, via the input parameter *TCRp*. This suggests a dependency between the level of TCR phosphorylation after vaccine input and the influence of this circadian rhythm, in such a way that there exists a threshold in *TCRp* below of which a circadian rhythm may not affect CD8^+^ T cell signaling. These results thus show that there is a potential complex relationship between the vaccine load presented to a CD8^+^ T cell and the molecular circadian clock.

Through a time-series analysis at the single-cell scale, we showed that only a circadian rhythm in the *S*_0_ state of the model produces a physiologically valid day-night variation and that such a circadian effect ceases when a vaccine is applied. This suggests that the physiological circadian rhythm acts during the baseline, pre-vaccination, state of a CD8^+^ T cell, ’priming’ them to respond to a vaccine. This result corroborates previously observed experimental results showing that CD8^+^ T cells are more primed for activation even prior to vaccination (Nobis et al., 2019b). Additionally, this finding is aligned with research in macrophages showing that the *Bmal*1 clock gene expression is reduced after an immune challenge via LPS (Curtis et al., 2015). To expand this analysis to other molecular players of CD8^+^ T cell signaling, the filtering properties of molecular players within the ZAP axis to the downstream LAT and CD28 axes were explored. Our results revealed that an *S*_0_ circadian rhythm within the ZAP axis is not transmitted to certain downstream signaling molecules within the CD28 axis, namely *TSC*1 *−* 2*, RhebGTP* and *mTORC*1. However, a circadian rhythmicity to their activity was restored by adding a circadian component to their own activation parameters, namely to *k*42*f*. Since a day-night variation was experimentally observed in *mTORC*1*^∗^* (Nobis et al., 2019b), our results suggest that this day-night variation may be due to independent circadian rhythms acting directly on *mTORC*1, rather than being due to the activity of an upstream circadian rhythm within the ZAP axis.

To produce a more physiological representation of CD8^+^ T cell responses to vaccination and how that is impacted by a circadian rhythm, we implemented a multi-dimensional population level modeling approach. This allowed us to explore the effect of a CD8^+^ T cell circadian rhythm on multiple molecular players simultaneously, through what we termed as a macro-bifurcation diagram. In these simulations, key circadian-dependent parameters were randomly sampled within their respective ranges, and produced three regions of activity with respect to vaccine load, defined by *TCRp*: insensitive, sensitive (macro-bistable) and elevated. Additionally, we demonstrated that adding additional rate activation parameters to the list of circadian-dependent parameters expanded the sensitive region of the macro-bifurcation diagram. This suggests that the increased variability in the number of circadian-dependent parameters increases the sensitivity of a population of CD8^+^ T cells to vaccination (by lowering the minimum *TCRp* needed for CD8^+^ T cells to be in the elevated steady state). That said, further analysis should be done to link this increased sensitivity to the circadian rhythm directly, by identifying which parameter sets bias T cells to the upper steady states, providing us a more complete understanding of the specifics of this interaction on a population level.

One limitation of this study is its assumption that changes in activated ZAP70 and mTOR translate to changes in CD8^+^ T cell activation. Our future work involves expanding this model to cover the downstream CD8^+^ gene regulatory outputs. This would expand the scope of the model and allow us to make direct predictions about cytokine production and subsequent proliferation of CD8^+^ T cells. This expansion would also assist in the experimental validation of the model, since outcomes would be closely linked to CD8^+^ T cell responses. Alternatively, a different reference model may be chosen as a starting point for this analysis using previously proposed mathematical models of T cell signaling Lipniacki et al. (2008); De Boer et al. (2001, 2003); Coombs et al. (2002); Fraņcois et al. (2013). That said, this study provided several novel insights into the effect of a circadian rhythm on CD8^+^ T cell signaling, in the context of vaccination, and serves a valid framework for assessing the influence of circadian rhythms in other models.

## Acknowledgments

This work was supported by the Natural Sciences and Engineering Council of Canada (http://www.nserc-crsng.gc.ca/index_eng.asp) discovery grant to A.K. and the Canadian Institutes of Health Research (https://cihr-irsc.gc.ca/e/193.html) grants program: MOP-119322 and PJT-186278 to N.C. N.B. was supported by the Canada Graduate Scholarships — Master’s program (https://www.nserc-crsng.gc.ca/ students-etudiants/pg-cs/cgsm-bescm_eng.asp).

## 6. Appendix

### 6.1. Model Parameterization

Here, we discuss the key reactions and framework of the model as well as note any modifications we made to accommodate CD8^+^ T cell signaling. Broadly, the model encompasses the immediate events following antigen presentation to a single T cell. The equations of the model are shown in Tables 1 and 2.

**Table 1:**
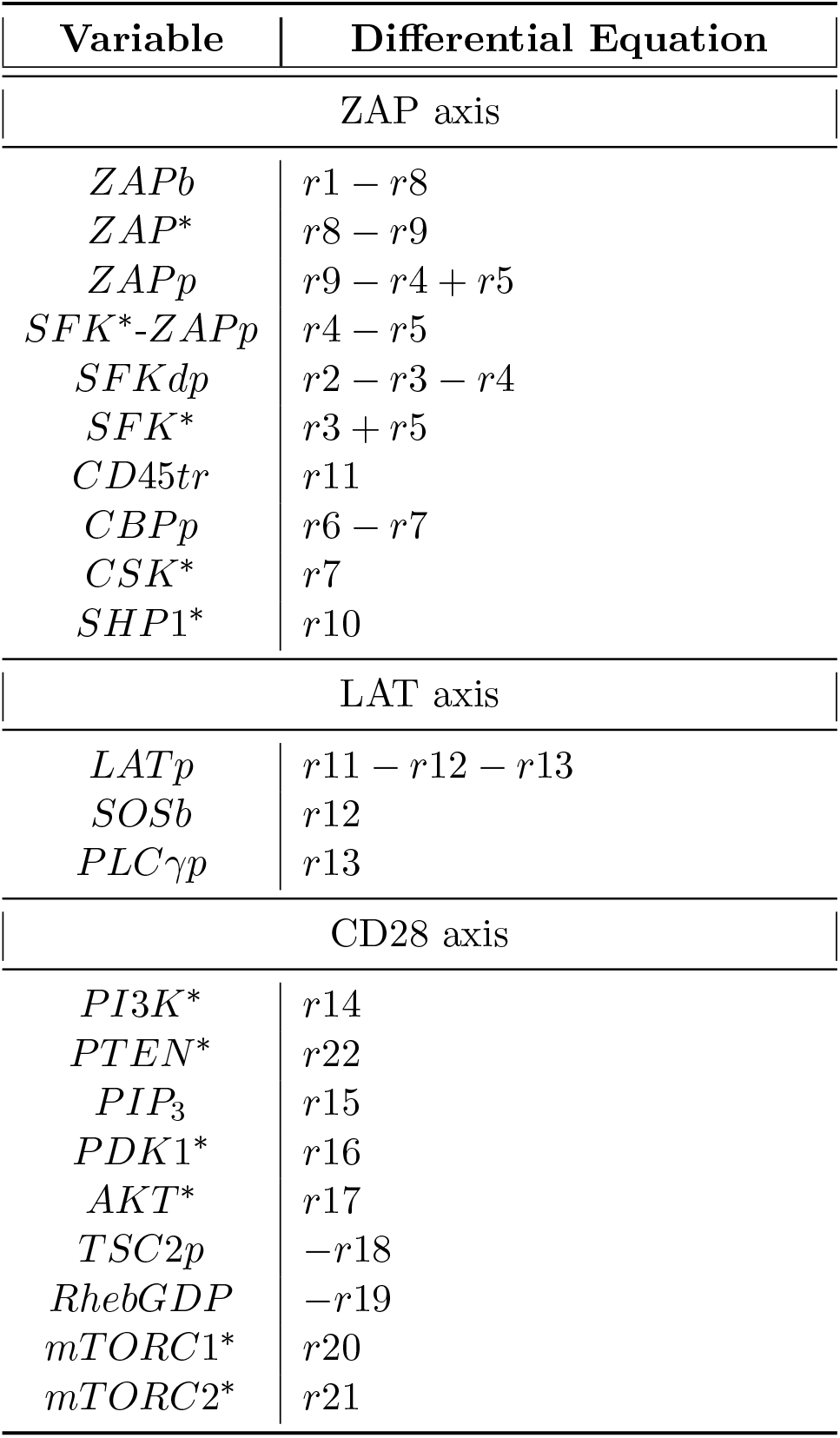
Differential equations of the various signaling molecules within the model. Equations adapted from Zheng et al. (2005) and Perley et al. (2014).

| Variable | Differential Equation |
| --- | --- |
| ZAP axis |  |
| $ZAPb$ | $r1 - r8$ |
| $ZAP^*$ | $r8 - r9$ |
| $ZAPp$ | $r9 - r4 + r5$ |
| $SFK^*-ZAPp$ | $r4 - r5$ |
| $SFKdp$ | $r2 - r3 - r4$ |
| $SFK^*$ | $r3 + r5$ |
| $CD45tr$ | $r11$ |
| $CBPp$ | $r6 - r7$ |
| $CSK^*$ | $r7$ |
| $SHP1^*$ | $r10$ |
| LAT axis |  |
| $LATp$ | $r11 - r12 - r13$ |
| $SOSb$ | $r12$ |
| $PLC\gamma p$ | $r13$ |
| CD28 axis |  |
| $PI3K^*$ | $r14$ |
| $PTEN^*$ | $r22$ |
| $PIP_3$ | $r15$ |
| $PDK1^*$ | $r16$ |
| $AKT^*$ | $r17$ |
| $TSC2p$ | $-r18$ |
| $RhebGDP$ | $-r19$ |
| $mTORC1^*$ | $r20$ |
| $mTORC2^*$ | $r21$ |

**Table 2:** The rate equations of the various signaling molecules within the model. Equations in the ZAP axis are multiplied by 60 to account for time-scale differences in parameter values. Model equations adapted from Zheng et al. (2005) and Perley et al. (2014).

| Rate Equation | Components |
| --- | --- |
| ZAP axis |  |
| $r1$ | $(k1f(2TCRp - ZAPb - ZAP^* - ZAPp - SFK^*-ZAPp)ZAP - k1r ZAPb) * 60$ |
| $r2$ | $(k2f CD45 SFK - k2r CSK^* SFKdp) * 60$ |
| $r3$ | $(k3f SFKdp - (k3r1 SHP1^* + k3r2) SFK^*) * 60$ |
| $r4$ | $(k4f SFK^*-ZAPp SFKdp - k4r SFK^*-ZAPp) * 60$ |
| $r5$ | $(k5f SFK^*-ZAPp - k5r SFK^* ZAPp) * 60$ |
| $r6$ | $((k6f1 SFK^* + k6f2)CBP - k6r CBPp) * 60$ |
| $r7$ | $(k7f CBP CSK - k7r CSK^*) * 60$ |
| $r8$ | $((k8f1 SFK^* + k8f2 SFK^*-ZAPp)ZAPb - (k8r1 SHP1^* + k8r2)ZAP^*) * 60$ |
| $r9$ | $((k9f1(ZAPp+ZAP^*+SFK^*-ZAPp)+k9f2 SFK^*+k9f3 SFK^*-ZAPp)ZAP^* - (k9r1 SHP1^* + k9r2)ZAPp) * 60$ |
| $r10$ | $(k10f SFK^* SHP1 - k10r SHP1^*) * 60$ |
| LAT axis |  |
| $r11$ | $k11f(ZAP^* + ZAPp + SFK^*-ZAPp)LAT - (k11r1 SHP1^* + k11r2)LATp$ |
| $r12$ | $k12f LATp GRB2SOS - k12r SOSb$ |
| $r13$ | $k13f LATp PLC\gamma - k13r PLC\gamma p$ |
| CD28 axis |  |
| $r14$ | $k14f CD28^*(ZAP^* + ZAPp + SFK^*-ZAPp)PI3K - k14r PI3K^*$ |
| $r15$ | $k15f PI3K^* PIP2 - (k15r1 PTEN^* + k15r2)PIP3$ |
| $r16$ | $k16f PIP3 PDK1 - k16r PDK1^*$ |
| $r17$ | $k17f1 PDK1^* + k17f2 mTORC2^* + k17f3 PDK1^* mTORC2^*)AKT - k17r AKT^*$ |
| $r18$ | $k18f AKT^* TSC12 - k18r TSCp$ |
| $r19$ | $k19f TSC12 RHEBgtp - k19r RHEBgdp$ |
| $r20$ | $k20f RHEBgtp mTORC1 - k20r mTORC1^*$ |
| $r21$ | $k21f PI3K^* mTORC2 - (k21r1 mTORC1^* + k21r2)mTORC2^*$ |
| $r22$ | $k22f PTEN - k22r(TCRp^{n22r}/(TCRp^{n22r} + kr22^{n22r}))PTEN^*$ |

**Table 3:** Parameter values of the ZAP axis; adapted from Zheng et al. (2005) and Perley et al. (2014).

| Parameter | Value | Units | Significance | Source |
| --- | --- | --- | --- | --- |
| $TCRp$ | 20000 | mol | Total number of phosphorylated receptor (model input) | Zheng et al. (2005) |
| $ZAP_{total}$ | 100000 | mol | Quantity of ZAP70 in cell | Zheng et al. (2005) |
| $SFK_{total}$ | 120000 | mol | Quantity of SFK in cell | Zheng et al. (2005) |
| $CD45_{total}$ | 100000 | mol | Quantity of CD45 in cell | Zheng et al. (2005) |
| $CBP_{total}$ | 50000 | mol | Quantity of CBP in cell | Zheng et al. (2005) |
| $CSK_{total}$ | 50000 | mol | Quantity of CSK in cell | Zheng et al. (2005) |
| $SHP1_{total}$ | 1000000 | mol | Quantity of SHP1 in cell | Zheng et al. (2005) |
| $k1f$ | $2 \cdot 10^{-5}$ | $(\text{mol} \cdot \text{s})^{-1}$ | Association rate of ZAP70-TCR binding | Zheng et al. (2005) |
| $k1r$ | $4.2 \cdot 10^{-2}$ | $\text{s}^{-1}$ | Dissociation rate of ZAP70-TCR binding | Zheng et al. (2005) |
| $k2f$ | $4 \cdot 10^{-7}$ | $(\text{mol} \cdot \text{s})^{-1}$ | Phosphorylation rate of SFK | Zheng et al. (2005) |
| $k2r$ | $4 \cdot 10^{-6}$ | $\text{s}^{-1}$ | Dephosphorylation rate of bound SFK | Zheng et al. (2005) |
| $k3f$ | $4 \cdot 10^{-3}$ | $\text{s}^{-1}$ | Phosphorylation rate of SFK by other kinases | Zheng et al. (2005) |
| $k3r1$ | $4 \cdot 10^{-6}$ | $(\text{mol} \cdot \text{s})^{-1}$ | Dephosphorylation rate of SFK* by CD45 or SHP1 | Zheng et al. (2005) |
| $k3r2$ | $8 \cdot 10^{-2}$ | $\text{s}^{-1}$ | Dephosphorylation rate of SFK* by other phosphatases | Zheng et al. (2005) |
| $k4f$ | $5 \cdot 10^{-6}$ | $(\text{mol} \cdot \text{s})^{-1}$ | Association rate of ZAPp to SFK via SH2 domains | Zheng et al. (2005) |
| $k4r$ | $8 \cdot 10^{-3}$ | $\text{s}^{-1}$ | Dissociation rate of ZAPp to SFK via SH2 domains | Zheng et al. (2005) |
| $k5f$ | $8 \cdot 10^{-2}$ | $\text{s}^{-1}$ | Dissociation rate of ZAPp to SFK via SH2 domains | Zheng et al. (2005) |
| $k5r$ | $1 \cdot 10^{-6}$ | $(\text{mol} \cdot \text{s})^{-1}$ | Association rate of ZAPp to SFK via SH2 domains | Zheng et al. (2005) |
| $k6f1$ | $8.33 \cdot 10^{-7}$ | $(\text{mol} \cdot \text{s})^{-1}$ | Phosphorylation rate of SFK by SFK | Zheng et al. (2005) |
| $k6f2$ | $1 \cdot 10^{-2}$ | $\text{s}^{-1}$ | Phosphorylation rate of SFK by other kinases | Zheng et al. (2005) |
| $k6r$ | $8 \cdot 10^{-7}$ | $(\text{mol} \cdot \text{s})^{-1}$ | Dephosphorylation rate of SFK | Zheng et al. (2005) |
| $k7f$ | $8.89 \cdot 10^{-7}$ | $(\text{mol} \cdot \text{s})^{-1}$ | Phosphorylation rate of CSK | Zheng et al. (2005) |
| $k7r$ | $4 \cdot 10^{-2}$ | $\text{s}^{-1}$ | Dephosphorylation rate of CSK | Zheng et al. (2005) |
| $k8f1$ | $4 \cdot 10^{-7}$ | $(\text{mol} \cdot \text{s})^{-1}$ | Tyrosine phosphorylation rate of bound ZAP by SFKdp | Manually derived |
| $k8f2$ | $2.5 \cdot 10^{-6}$ | $(\text{mol} \cdot \text{s})^{-1}$ | Tyrosine phosphorylation rate of bound ZAP by SFK* | Zheng et al. (2005) |
| $k8r1$ | $4 \cdot 10^{-6}$ | $(\text{mol} \cdot \text{s})^{-1}$ | Dephosphorylation rate of ZAP* by SHP1 | Zheng et al. (2005) |
| $k8r2$ | $8 \cdot 10^{-2}$ | $\text{s}^{-1}$ | Dephosphorylation rate of ZAP* by other phosphatases | Zheng et al. (2005) |
| $k9f1$ | $4.44 \cdot 10^{-6}$ | $(\text{mol} \cdot \text{s})^{-1}$ | Tyrosine phosphorylation rate of ZAP* by activated ZAP | Zheng et al. (2005) |
| $k9f2$ | $8.33 \cdot 10^{-7}$ | $(\text{mol} \cdot \text{s})^{-1}$ | Tyrosine phosphorylation rate of ZAP* by SFKdp | Zheng et al. (2005) |
| $k9f3$ | $2.5 \cdot 10^{-6}$ | $(\text{mol} \cdot \text{s})^{-1}$ | Tyrosine phosphorylation rate of ZAP* by SFK* | Zheng et al. (2005) |
| $k9r1$ | $4 \cdot 10^{-6}$ | $(\text{mol} \cdot \text{s})^{-1}$ | Dephosphorylation rate of ZAPp by SHP1 | Zheng et al. (2005) |
| $k9r2$ | $8 \cdot 10^{-2}$ | $\text{s}^{-1}$ | Dephosphorylation rate of ZAPp by other phosphatases | Zheng et al. (2005) |
| $k10f$ | $8.64 \cdot 10^{-10}$ | $(\text{mol} \cdot \text{s})^{-1}$ | Activation rate of SHP1 | Zheng et al. (2005) |
| $k10r$ | $4 \cdot 10^{-4}$ | $\text{s}^{-1}$ | Deactivation rate of SHP1 | Zheng et al. (2005) |
| $k11f$ | $4 \cdot 10^{-8}$ | $(\text{mol} \cdot \text{s})^{-1}$ | Activation rate of CD45 | Zheng et al. (2005) |
| $k11r$ | $2 \cdot 10^{-4}$ | $\text{s}^{-1}$ | Deactivation rate of CD45 | Zheng et al. (2005) |

**Table 4:** Parameter values of the LAT axis; adapted from Zheng et al.Zheng et al. (2005) and Perley et al.Perley et al. (2014).

| Parameter | Value | Units | Significance | Source |
| --- | --- | --- | --- | --- |
| $LAT_{total}$ | 50000 | mol | Quantity of LAT in cell | Perley et al. (2014) |
| $PLC\gamma_{total}$ | 50000 | mol | Quantity of PLCγ in cell | Perley et al. (2014) |
| $GRB2SOS_{total}$ | 50000 | mol | Quantity of GRB2SOS in cell | Perley et al. (2014) |
| $PIP2_{total}$ | 10000000 | mol | Quantity of PIP2 in cell | Perley et al. (2014) |
| $k12f$ | $3 \cdot 10^{-4}$ | $(\text{mol} \cdot \text{min})^{-1}$ | Phosphorylation rate of LAT by activated ZAP | Perley et al. (2014) |
| $k12r1$ | $2 \cdot 10^{-3}$ | $(\text{mol} \cdot \text{min})^{-1}$ | Dephosphorylation rate of LAT by activated SHP1 | Perley et al. (2014) |
| $k_{12r2}$ | $9 \cdot 10$ | $\text{min}^{-1}$ | Dephosphorylation rate of LAT by other phosphatases | Perley et al. (2014) |
| $k_{13f}$ | $3 \cdot 10^{-3}$ | $(\text{mol} \cdot \text{min})^{-1}$ | Association rate of PLCγ to LAT complex | Perley et al. (2014) |
| $k_{13r}$ | $3 \cdot 10^2$ | $\text{min}^{-1}$ | Dissociation rate of PLCγ from LAT complex | Perley et al. (2014) |
| $k_{14f}$ | $3 \cdot 10^{-5}$ | $(\text{mol} \cdot \text{min})^{-1}$ | Association rate of SOSb to LAT complex | Perley et al. (2014) |
| $k_{14r}$ | $6 \cdot 10^{-2}$ | $\text{min}^{-1}$ | Dissociation rate of SOSb from LAT complex | Perley et al. (2014) |

**Table 5:** Parameter values of the CD28 axis; adapted from Zheng et al.Zheng et al. (2005) and Perley et al.Perley et al. (2014).

| Parameter | Value | Units | Significance | Source |
| --- | --- | --- | --- | --- |
| $CD28^*$ | 20000 | mol | Total number of activated co-receptor (input signal) | Manually derived |
| $PI3K_{total}$ | 10000 | mol | Quantity of PI3K in cell | Perley et al. (2014) |
| $PDK1_{total}$ | 10000 | mol | Quantity of PDK1 in cell | Perley et al. (2014) |
| $AKT_{total}$ | 10000 | mol | Quantity of AKT in cell | Perley et al. (2014) |
| $TSC_{total}$ | 10000 | mol | Quantity of TSC in cell | Perley et al. (2014) |
| $RHEB_{total}$ | 10000 | mol | Quantity of RHEB in cell | Perley et al. (2014) |
| $mTORC1_{total}$ | 10000 | mol | Quantity of mTORC1 in cell | Perley et al. (2014) |
| $mTORC2_{total}$ | 10000 | mol | Quantity of mTORC2 in cell | Perley et al. (2014) |
| $PTEN_{total}$ | 10000 | mol | Quantity of PTEN in cell | Perley et al. (2014) |
| $k_{15f}$ | $3 \cdot 10^{-9}$ | $(\text{mol}^2 \cdot \text{min})^{-1}$ | Activation rate of PI3K by activated CD28 | Perley et al. (2014) |
| $k_{15r}$ | $3 \cdot 10$ | $\text{min}^{-1}$ | Dissociation rate of PI3K | Perley et al. (2014) |
| $k_{16f}$ | 3 | $(\text{mol} \cdot \text{min})^{-1}$ | Phosphorylation rate of PIP2 by PI3K | Perley et al. (2014) |
| $k_{16r1}$ | $3 \cdot 10$ | $(\text{mol} \cdot \text{min})^{-1}$ | Dephosphorylation rate of PIP3 by PTEN | Perley et al. (2014) |
| $k_{16r2}$ | $1 \cdot 10^{-10}$ | $\text{min}^{-1}$ | Dephosphorylation rate of PIP3 by other phosphatases | Perley et al. (2014) |
| $k_{17f}$ | $3 \cdot 10^{-5}$ | $(\text{mol}^2 \cdot \text{min})^{-1}$ | Activation rate of PDK1 by PIP3 | Perley et al. (2014) |
| $k_{17r}$ | $3 \cdot 10$ | $\text{min}^{-1}$ | Dissociation rate of PDK1* | Perley et al. (2014) |
| $k_{18f1}$ | $1 \cdot 10^{-8}$ | $(\text{mol} \cdot \text{min})^{-1}$ | Phosphorylation rate of AKT by PDK1* | Manually derived |
| $k_{18f2}$ | $1 \cdot 10^{-11}$ | $(\text{mol} \cdot \text{min})^{-1}$ | Phosphorylation rate of AKT by mTORC2* | Manually derived |
| $k_{18f3}$ | $1 \cdot 10^{-8}$ | $(\text{mol}^2 \cdot \text{min})^{-1}$ | Phosphorylation rate of AKT by PDK1* and mTORC2* | Perley et al. (2014) |
| $k_{18r}$ | $1 \cdot 10^{-1}$ | $\text{min}^{-1}$ | Dephosphorylation rate of AKT | Manually derived |
| $k_{19f}$ | $1 \cdot 10$ | $(\text{mol} \cdot \text{min})^{-1}$ | Phosphorylation of TSC2 subunit of TSC1-2 complex | Perley et al. (2014) |
| $k_{19r}$ | 1 | $\text{min}^{-1}$ | Association of TSC2 to TSC1 | Perley et al. (2014) |
| $k_{20f}$ | $1 \cdot 10^{-4}$ | $(\text{mol} \cdot \text{min})^{-1}$ | Activation rate of RHEBgtp | Perley et al. (2014) |
| $k_{20r}$ | $1 \cdot 10^{-1}$ | $\text{min}^{-1}$ | Dissociation rate of RHEBgtp | Manually derived |
| $k_{21f}$ | $1 \cdot 10^{-4}$ | $(\text{mol} \cdot \text{min})^{-1}$ | Activation rate of mTORC1 by RHEBGTP | Perley et al. (2014) |
| $k_{21r}$ | 1 | $\text{min}^{-1}$ | Dissociation rate of mTORC1 by RHEBGTP | Perley et al. (2014) |
| $k_{22f}$ | $3 \cdot 10^{-3}$ | $(\text{mol} \cdot \text{min})^{-1}$ | Activation rate of mTORC2 by PI3K* | Perley et al. (2014) |
| $k_{22r1}$ | $3 \cdot 10^{-3}$ | $(\text{mol} \cdot \text{min})^{-1}$ | Dissociation rate of mTORC2 by mTORC1* | Perley et al. (2014) |
| $k_{22r2}$ | 3 | $\text{min}^{-1}$ | Dissociation rate of mTORC2 by other phosphatases | Perley et al. (2014) |
| $k_{23f}$ | 1 | $(\text{mol} \cdot \text{min})^{-1}$ | Activation rate of PTEN | Perley et al. (2014) |
| $k_{23r}$ | $2 \cdot 10$ | $\text{min}^{-1}$ | Deactivation rate of PTEN* by TCR complex | Perley et al. (2014) |
| $n_{23r}$ | $1 \cdot 10$ | | PTEN* hill coefficient of deactivation | Perley et al. (2014) |
| $kr_{23}$ | $0.75 \cdot TCRp$ | mol | Quantity of TCRp producing half maximum PTEN* deactivation | Perley et al. (2014) |

### 6.2. Oscillatory dynamics of the model

With the parameterization of the model described in Zheng et al. (2005) and Perley et al. (2014), the model is able to produce intrinsic oscillatory signals, shown in Fig. S1(A). For instance, setting *TCRp* = 20000 mol and *k*7*f* = 4.89 *·* 10*^−^*^7^ (mol *·* s)*^−^*^1^ and plotting the one-parameter bifurcation of *ZAPp* with respect to *k*10*f* (the rate constant for *SHP* 1 activation), as shown in Fig. S1(B), produces a diagram that effectively eliminates the bistable behavior of the model and produces an oscillatory regime defined by the envelopes of stable limit cycles that originate from saddle-node of periodic orbits (SNP) and terminate at a saddle-node on an invariant circle (SNIC) bifurcation (formed by SN2 and a homoclinic bifurcation HM). The stable envelopes of limit cycles produce periodic solutions with periodic spiking in *ZAP_active_* similar to that shown in Fig. S1(A).

This spiking pattern in *ZAP_active_* has not been observed experimentally, indicating that this is a non-viable parameter regime of the model. Physiologically, since *ZAP* 70 remains bound to the TCR throughout antigen presentation, this spiking pattern suggests that *ZAP* 70 is repeatedly getting phosphorylated and subsequently dephosphorylated in the time scale of seconds, all while remaining bound to the phosphorylated *TCR* (via the non-zero *TCRp* parameter). This result contradicts the literature showing a continuous level of phosphorylated *ZAP* 70 throughout antigen stimulation (Houtman et al. (2005); An et al. (2019)). The lack of evidence showing a spiking *ZAP* 70 pattern as well as the contrast to experimentally observed *ZAP* 70 binding/unbinding kinetics leads us to designate this regime as an invalid result of the model, as this behaviour is unlikely to exist. To resolve this issue, we have developed a systematic framework to eliminate this oscillatory regime while maintaining the other model regimes. This is the focus of the following section.

### 6.3. Eliminating the intrinsic oscillatory regime

This approach is adapted from the Dynamics Visualisation based Parallel Coordinates (DYVIPAC) framework by Nguyen et al. (2015). Broadly, it involves a population-based approach to uncover system’s dynamics. The *m* parameters whose effects on model dynamics are the main focus of the analysis are first selected, and a range of values for each parameter is set. A sample of *n* model realizations (i.e. simulated steady state responses) is then created, each with a randomly selected parameterization from within predetermined ranges. These realizations are integrated starting from the same initial conditions, and their steady state solutions are classified according to outcomes. A parallel coordinates plot is then created linking each realization of the model to its respective steady state classification. This allows us to obtain a multi-dimensional view of model dynamics, and allows us to reliably identify parameter regimes where the oscillatory solution exists. By limiting our parameter sampling ranges to exclude such parameter regimes, the oscillatory solutions can be excluded from our model. This approach is explored in more detail in the following sections.

**Figure S1:**
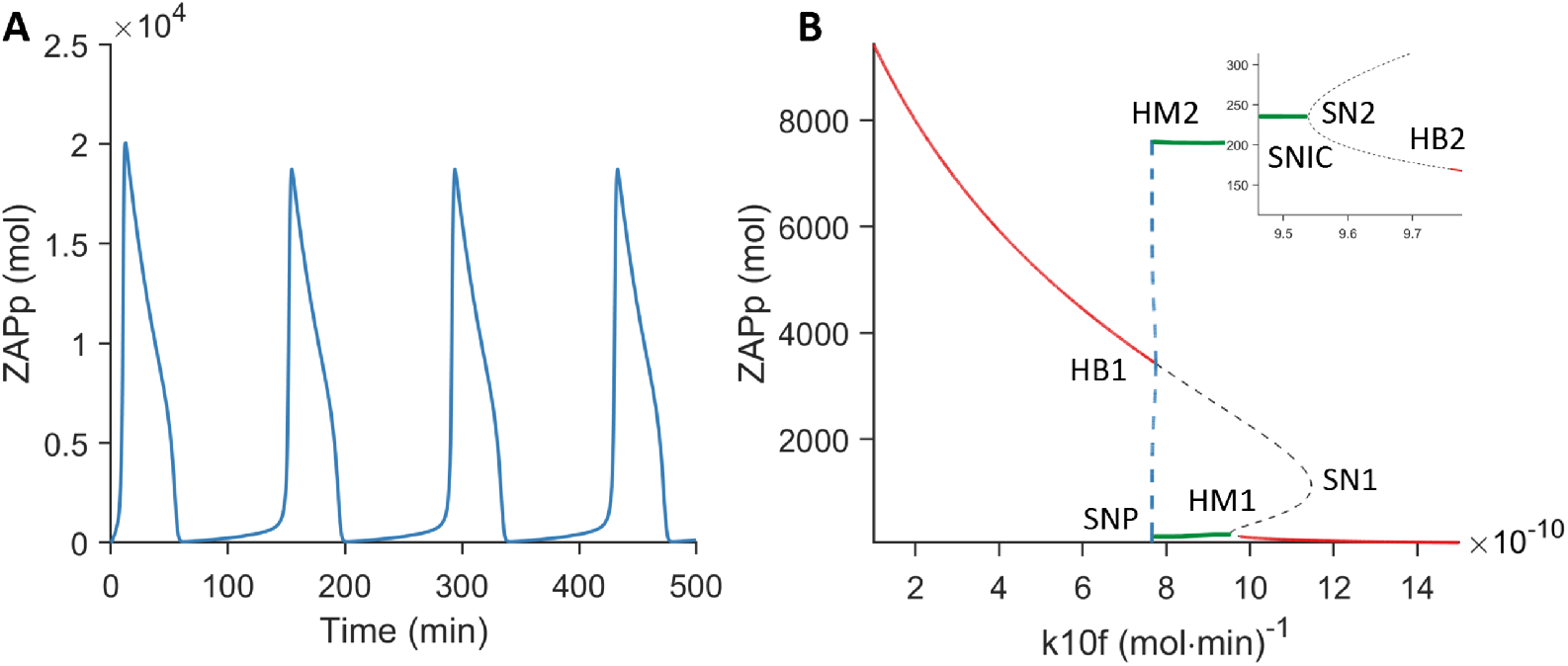
(A) Time series simulation of an oscillatory solution of *ZAP_active_* obtained by setting *TCRp* = 20000 mol*, k*7*f* = 4.89 *·* 10*^−^*^7^ (mol *·* s)*^−^*^1^ and *k*10*f* = 9 *·* 10*^−^*^10^ (mol *·* s)*^−^*^1^. (B) One-parameter bifurcation of phosphorylated ZAP (*ZAPp*) with respect to *k*10*f* when *TCRp* = 20000 mol and *k*7*f* = 4.89 *·* 10*^−^*^7^ (mol *·* s)*^−^*^1^. Red solid (gray dashed) lines represent stable (unstable) branches of equilibria, while green solid (blue dashed) lines represent envelopes of stable (unstable) limit cycles. HB1 and HB2: Hopf bifurcations, HM1 and HM2: homoclinic bifurcations, SN1 and SN2: saddle-node bifurcations, SNP: saddle-node of periodic orbits bifurcation, SNIC: saddle-node on an invariant circle bifurcation formed by SN2 and HM1 (see inset for magnification). Note that the periodic continuation (green solid line) was superimposed afterwards due to a lack of convergence.

#### 6.3.1. Network model and parameter selection

The first step of this framework is to identify the parameters of interest, or more specifically those that we vary in this study. These parameters are *TCRp*, *k*10*f* (which represents SHP1 activation) and *k*8*f* 1 (which represents ZAP activation); the latter is selected since it allows us to more easily eliminate the oscillatory regime. The number of parameters defines the dimensionality *m* of this approach (i.e., *m* = 3).

#### 6.3.2. Parameter sampling

With the dimensionality *m* of the framework determined through parameter selection, an appropriate sampling range of these parameters now needs to be set. Each realization of the model will have a randomly selected value, chosen from within this range. Since we are interested in the effects of a CD8^+^ T cell circadian rhythm, this parameter range should include the degree of modulation due to this circadian rhythm. This range is therefore defined by

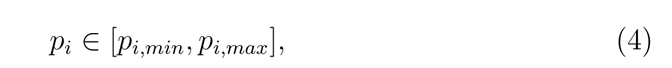

where *p_i_*is the the value of a parameter *i* randomly sampled from within a set range. This *p_i_*, when sampled over a sufficiently large *n* realizations, demonstrates all the possible values this parameter can have when subjected to a circadian rhythm. *p_i,_*_0_ denotes the default parameter value, and *p_i,min_*and *p_i,max_* denote the minimum and maximum endpoints from which this parameter can be sampled from.

#### 6.3.3. Setting a physiological p_i_ range

To impose a physiologically-relevant limit for the influence of this circadian rhythm, we note that in CD8^+^ T cell signaling, it was shown that IFN*γ*, along with other CD8^+^ T cell cytokines, approximately vary within a [0.5 *× p*_0_, 1.5 *× p*_0_] range depending on the time of initial vaccination (Fortier et al., 2011; Nobis et al., 2019b). We will therefore use this range to describe the influence of a circadian rhythm on CD8^+^ T cell signaling. To produce this *p* range, we choose the range of a viable circadian coefficient *α* to be *α ∈* [0, 0.5]. It should be noted that this only defines the maximum range of *p*; some realizations, *k* = 1, 2*, . . . , n*, of the model may have a parameter subjected to a weaker circadian influence (defined by an *α <* 0.5).

It is also important to note that the above range assumes that a [0.5 *× p*_0_, 1.5 *× p*_0_] variation in cytokine production can be traced linearly to a variation of a similar magnitude in ZAP signaling. While we show in this study that the axes downstream of ZAP do not contribute significantly to model dynamics, we acknowledge that there may be additional factors contributing to the variation experimentally seen in IFN*γ* and other cytokines. Regardless, this assumption is made in this study to facilitate our analysis and selection of a viable parameter range.

### 6.4. Analysis of dynamic behaviour

With *n* model realizations selected from parameters within physiological ranges, each individual realization, *k* = 1, 2*, . . . , n*, can be integrated and the steady-state solution associated with each one analyzed and classified according to its properties. Whereas DYVIPAC uses model eigenvalues to calculate stability (Nguyen et al., 2015), our approach classifies solutions based on their steady state behaviour. Namely, we classify solutions as either elevated (i.e., corresponding to those with a higher than 5% active ZAP concentration), oscillatory (i.e., corresponding to those with a slope greater than a manually selected threshold of 1) or inactive (i.e., corresponding to those fulfilling neither of the previous criteria). We have found that performing the classification in this manner is effective due to two important dynamic features of the model, including: (i) its fast dynamics, allowing it to reach steady state relatively quickly, and (ii) the relaxation oscillation-like behaviour it exhibits within the oscillatory regime with its rapid up and down-stroke oscillations that facilitate our ability to classify it according to its slope.

With this, we are left with a matrix of *n* model realizations, each with its unique parameterization of the *m* circadian-affected parameters, and with a label designating the classification of each solution *k* = 1, 2*, . . . , n* based on the properties of its steady-state dynamics.

### 6.5. Parallel coordinate plot visualization

This dataset is visualized using a parallel coordinate plot, which displays how these different parameter values can influence the dynamics of the model. Each model realization is linked to its respective steady state dynamics according to the values of its parameters of interest. Such visualization is shown in Figure S2(A), where a multi-dimensional analysis of three parameters, including *TCRp*, *k*10*f* and *k*8*f* 1, is conducted. In this analysis, *n* = 1000 model realizations, where each realization is represented individually by a colored segmented line linking the parameter values generating this realization. The parameter values of each realization lie above the ticks on the x-axis representing each parameter, and are connected by a segmented line color-coded according to the stability property of that particular model realization (i.e., the stability of the steady-state solution associated with that realization). Three colors are used, blue, red and yellow representing inactive, elevated and oscillatory solutions, respectively.

The parameter values are scaled between [0, 1] according to their respective maximum and minimum sample values. This normalization is generated using the equation

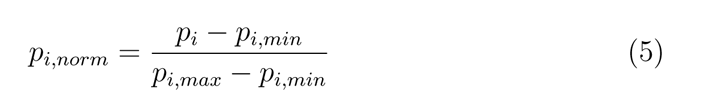

where, *p_i,norm_* is the normalized parameter value between [0, 1] of a specific parameter of interest, *p_i_* is the parameter value of a particular model realization, and *p_i,max_* and *p_i,min_* are the respective maximum and minimum parameter values across all *n* realizations. This visualization allows us to obtain a broader picture of model dynamics and how these dynamics can be shaped by different parameter combinations.

#### 6.5.1. Eliminating the oscillatory regime

Using the parallel coordinate plot, the oscillatory regime can be extracted from the original scaled range of an indicator parameter *p_i_* by removing the continuous interval within this range associated with the oscillatory behaviour (i.e., the interval that has segmented yellow lines). The indicator parameter *p_i_* is then re-scaled according to the remaining interval and sampling is then restricted to this constrained range (which by design excludes oscillatory solutions). This is done in Fig. S2(B), where the elevated and inactive solutions from the previously presented parallel coordinate plot are hidden, to better visualize the oscillatory regimes.

Using this approach indicates that the oscillatory solutions are confined to a normalized *k*8*f* 1 range of [0.23, 1]. Thus, by only sampling from *k*8*f* 1 *∈* [0, 0.23], the oscillatory regime can be completely avoided.

Using this threshold of *k*8*f* 1*_max_* = 0.23 (mol *·* s)*^−^*^1^, we reverse the previously described normalization process, to obtain the raw maximum value of *k*8*f* 1*_max_* = 6.12 (mol *·* s)*^−^*^1^. To obtain the new *p_i,_*_0_ for *k*8*f* 1, the trivial calculation of *p_i,_*_0_ = *k*8*f* 1*_max_/*1.5 is performed. The preceding calculation is repeated to find *k*8*f* 1*_min_*, thus producing a new range *k*8*f* 1 *∈* [2.72, 6.12] (mol *·* s)*^−^*^1^. Applying parallel coordinate plot on this new regime now only produces two outcomes: the inactive and elevated states, as shown in Figure S2(C). Through this new *k*8*f* 1 parameterization, we observe that the oscillatory regime is indeed eliminated, while still preserving both the inactive and elevated model solution. Thus, this new model parameterization can safely allow us to simulate the effects of a circadian rhythm on CD8^+^ T cell signaling impacting directly *k*10*f* and *k*8*f* 1 without worrying about intrinsic oscillatory dynamics that could be produced by the model; this is because we are certain that any combination of these three parameters under this parameterization considered in Figure S2(C) would not produce the non-physiological oscillatory solutions. This constrained parameter regime of *k*8*f* 1 is used in both our single-cell and population-level modeling simulations to avoid the emergence of oscillatory solutions.

**Figure S2:**
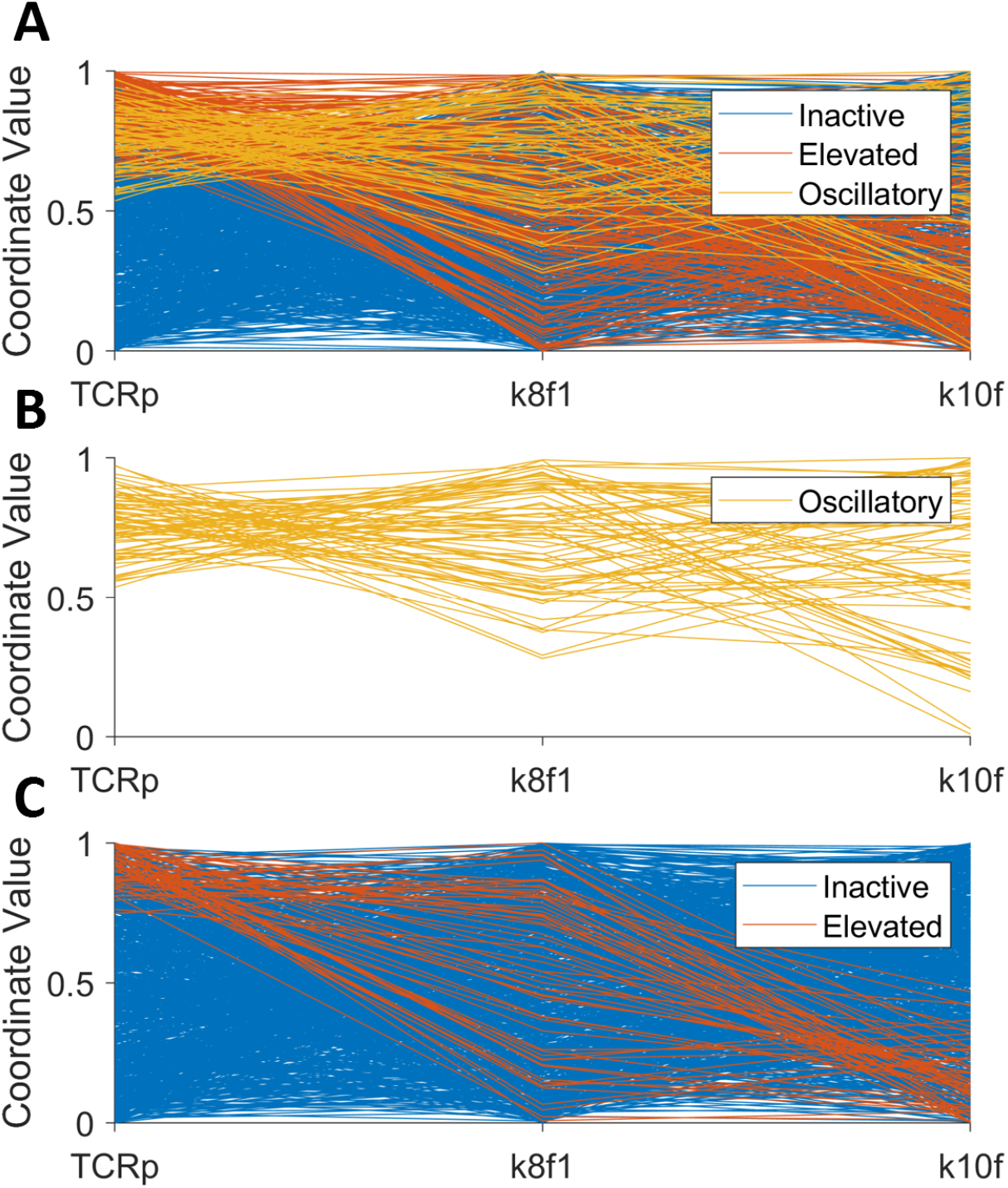
Parallel coordinate plot of *TCRp, k*8*f* 1*, k*10*f* . (A) The three parameters are displayed on the x-axis and their normalized parameter values are displayed on the yaxis. Each segmented colored line represents a given model realization (i.e., steady state response) sampled randomly from within a pre-determined range. The segmented lines are color-coded according to the type of steady state obtained: blue for inactive, red for elevated and yellow for oscillatory solution. (B) The isolated oscillatory solutions (yellow) from (A). Range of parameter values for (A) & (B): *TCRp ∈* [0, 30000], *k*8*f* 1 *∈* [4.17, 12.50], *k*10*f ∈* [4.32, 12.95]. (C) The analysis repeated with a constrained parameter range for *k*8*f* 1. Notice the lack of yellow oscillatory solutions. Range of parameter values for (C): *TCRp ∈* [0, 30000], *k*8*f* 1 *∈* [2, 6], *k*10*f ∈* [4.32, 12.95].

## References

1. An, W., Park, K., Reyes, A., Kwieraga, A., Sabri, N., Frey, A., Chronopoulou, E., Savas-Carstens, J., Soper, D., Elia, J., et al., 2019. The phosphorylation kinetics of the key regulators in T cell receptor signaling pathway.

2. Bai, Z., Zhou, Y., 2012. Dynamics of a viral infection model with delayed CTL response and immune circadian rhythm. Chaos, Solitons and Fractals 45, 1133–1139. URL: https://www.sciencedirect.com/science/article/pii/S0960077912001245, 10.1016/j.chaos.2012.06.001.

3. Boivin, D.B., James, F.O., Wu, A., Cho-Park, P.F., Xiong, H., Sun, Z.S., 2003. Circadian clock genes oscillate in human peripheral blood mononuclear cells. Blood 102, 4143–4145.

4. Bollinger, T., Leutz, A., Leliavski, A., Skrum, L., Kovac, J., Bonacina, L., Benedict, C., Lange, T., Westermann, J., Oster, H., et al., 2011. Circadian clocks in mouse and human cd4+ t cells. PloS one 6, e29801.

5. de Bree, L.C.J., Mourits, V.P., Koeken, V.A., Moorlag, S.J., Janssen, R., Folkman, L., Barreca, D., Krausgruber, T., Fife-Gernedl, V., Novakovic, B., et al., 2020. Circadian rhythm influences induction of trained immunity by bcg vaccination. The Journal of clinical investigation 130, 5603–5617.

6. Cermakian, N., Stegeman, S.K., Tekade, K., Labrecque, N., 2021. Circadian rhythms in adaptive immunity and vaccination, in: Seminars in Immunopathology, Springer. pp. 1–15.

7. Chi, H., 2012. Regulation and function of mtor signalling in t cell fate decisions. Nature reviews immunology 12, 325–338.

8. Coombs, D., Dushek, O., Merwe, P., 2011. A review of mathematical models for T cell receptor triggering and antigen discrimination. Mathematical models and immune cell biology , 25–45.

9. Coombs, D., Kalergis, A.M., Nathenson, S.G., Wofsy, C., Goldstein, B., 2002. Activated tcrs remain marked for internalization after dissociation from pmhc. Nature Immunology 3, 926–931. doi:10.1038/ni838.

10. Cuesta, M., Boudreau, P., Cermakian, N., Boivin, D.B., 2017. Rapid resetting of human peripheral clocks by phototherapy during simulated night shift work. Scientific reports 7, 1–11.

11. Curtis, A.M., Fagundes, C.T., Yang, G., Palsson-McDermott, E.M., Wochal, P., McGettrick, A.F., Foley, N.H., Early, J.O., Chen, L., Zhang, H., et al., 2015. Circadian control of innate immunity in macrophages by miR-155 targeting Bmal1. Proceedings of the National Academy of Sciences 112, 7231–7236.

12. De Boer, R.J., Homann, D., Perelson, A.S., 2003. Different dynamics of CD4+ and CD8+ T cell responses during and after acute lymphocytic choriomeningitis virus infection. The Journal of Immunology 171, 3928– 3935.

13. De Boer, R.J., Oprea, M., Antia, R., Murali-Krishna, K., Ahmed, R., Perelson, A.S., 2001. Recruitment times, proliferation, and apoptosis rates during the CD8+ T-cell response to lymphocytic choriomeningitis virus. Journal of virology 75, 10663–10669.

14. Dibner, C., Schibler, U., Albrecht, U., 2010. The mammalian circadian timing system: organization and coordination of central and peripheral clocks. Annual review of physiology 72, 517–549.

15. Dimitrov, S., Benedict, C., Heutling, D., Westermann, J., Born, J., Lange, T., 2009. Cortisol and epinephrine control opposing circadian rhythms in t cell subsets. Blood, the Journal of the American Society of Hematology 113, 5134–5143.

16. Elliott, A.L., Mills, J., Minors, D., Waterhouse, J., 1972. The effect of real and simulated time-zone shifts upon the circadian rhythms of body temperature, plasma 11-hydroxycorticosteroids, and renal excretion in human subjects. The Journal of Physiology 221, 227–257.

17. Esquifino, A.I., Selgas, L., Arce, A., Maggiore, V.D., Cardinali, D.P., 1996. Twenty-four-hour rhythms in immune responses in rat submaxillary lymph nodes and spleen: effect of cyclosporine. Brain Behav Immun 10, 92–102. doi:10.1006/brbi.1996.0010.

18. Fan, A., Wang, K., 2010. A viral infection model with immune circadian rhythms. Applied Mathematics and Computation 215, 3369–3374. URL: https://www.sciencedirect.com/science/article/pii/S0096300309009333, 10.1016/j.amc.2009. 10.028.

19. Fortier, E.E., Rooney, J., Dardente, H., Hardy, M.P., Labrecque, N., Cermakian, N., 2011. Circadian variation of the response of T cells to antigen. J Immunol 187, 6291–300. doi:10.4049/jimmunol.1004030.

20. Fraņcois, P., Voisinne, G., Siggia, E.D., Altan-Bonnet, G., Vergassola, M., 2013. Phenotypic model for early t-cell activation displaying sensitivity, specificity, and antagonism. Proceedings of the National Academy of Sciences 110, E888–E897.

21. de Goede, P., Wefers, J., Brombacher, E.C., Schrauwen, P., Kalsbeek, A., 2018. Circadian rhythms in mitochondrial respiration. J Mol Endocrinol 60, R115–r130. doi:10.1530/jme-17-0196.

22. Halberg, F., Johnson, E.A., Brown, B.W., Bittner, J.J., 1960. Susceptibility rhythm to e. coli endotoxin and bioassay. Proceedings of the Society for Experimental Biology and Medicine 103, 142–144.

23. Hazan, G., Duek, O.A., Alapi, H., Mok, H., Ganninger, A., Ostendorf, E., Gierasch, C., Chodick, G., Greenberg, D., Haspel, J.A., et al., 2023. Biological rhythms in covid-19 vaccine effectiveness in an observational cohort study of 1.5 million patients. The Journal of clinical investigation 133.

24. Hickey, D., Kirkland, J., Lucas, S., Lye, M., 1984. Analysis of circadian rhythms by fitting a least squares sine curve. Computers in biology and medicine 14, 217–223.

25. Hosking, M.P., Flynn, C.T., Whitton, J.L., 2014. Antigen-specific naive cd8+ t cells produce a single pulse of ifn-*γ* in vivo within hours of infection, but without antiviral effect. The Journal of Immunology 193, 1873–1885.

26. Houtman, J.C., Houghtling, R.A., Barda-Saad, M., Toda, Y., Samelson, L.E., 2005. Early phosphorylation kinetics of proteins involved in proximal TCRmediated signaling pathways. The Journal of Immunology 175, 2449–2458.

27. Jerigova, V., Zeman, M., Okuliarova, M., 2022. Circadian disruption and consequences on innate immunity and inflammatory response. International Journal of Molecular Sciences 23, 13722.

28. Ji, Y., Min, L., Zheng, Y., Su, Y., 2010. A viral infection model with periodic immune response and nonlinear CTL response. Mathematics and Computers in Simulation 80, 2309–2316. URL: https://www.sciencedirect.com/science/article/pii/S0378475410001539, 10.1016/j.matcom.2010.04.029.

29. Kawate, T., Abo, T., Hinuma, S., Kumagai, K., 1981. Studies of the bioperiodicity of the immune response. II. Co-variations of murine T and B cells and a role of corticosteroid. The Journal of Immunology 126, 1364. URL: http://www.jimmunol.org/content/126/4/1364.abstract.

30. Keller, M., Mazuch, J., Abraham, U., Eom, G.D., Herzog, E.D., Volk, H.D., Kramer, A., Maier, B., 2009. A circadian clock in macrophages controls inflammatory immune responses. Proceedings of the National Academy of Sciences 106, 21407–21412. URL: https://www.pnas.org/doi/abs/10.1073/pnas.0906361106, doi:doi:10.1073/pnas.0906361106.

31. Labrecque, N., Cermakian, N., 2015. Circadian clocks in the immune system. J Biol Rhythms 30, 277–90. doi:10.1177/0748730415577723.

32. Lipniacki, T., Hat, B., Faeder, J.R., Hlavacek, W.S., 2008. Stochastic effects and bistability in T cell receptor signaling. Journal of Theoretical Biology 254, 110–122. URL: https://www.sciencedirect.com/science/article/pii/S0022519308002208, 10.1016/j.jtbi.2008.05.001.

33. Liu, J., Mankani, G., Shi, X., Meyer, M., Cunningham-Runddles, S., Ma, X., Sun, Z.S., 2006. The circadian clock period 2 gene regulates gamma interferon production of nk cells in host response to lipopolysaccharideinduced endotoxic shock. Infection and immunity 74, 4750–4756.

34. Long, J.E., Drayson, M.T., Taylor, A.E., Toellner, K.M., Lord, J.M., Phillips, A.C., 2016. Morning vaccination enhances antibody response over afternoon vaccination: a cluster-randomised trial. Vaccine 34, 2679–2685.

35. Marino, S., Hogue, I.B., Ray, C.J., Kirschner, D.E., 2008. A methodology for performing global uncertainty and sensitivity analysis in systems biology. Journal of theoretical biology 254, 178–196.

36. Mørch, A.M., Bálint, S^̌^., Santos, A.M., Davis, S.J., Dustin, M.L., 2020. Coreceptors and tcr signaling–the strong and the weak of it. Frontiers in cell and developmental biology 8, 597627.

37. Naitoh, P., Englund, C.E., Ryman, D.H., 1985. Circadian rhythms determined by cosine curve fitting: analysis of continuous work and sleep-loss data. Behavior Research Methods, Instruments, & Computers 17, 630–641.

38. Nguyen, K.D., Fentress, S.J., Qiu, Y., Yun, K., Cox, J.S., Chawla, A., 2013. Circadian gene bmal1 regulates diurnal oscillations of ly6chi inflammatory monocytes. Science 341, 1483–1488.

39. Nguyen, L.K., Degasperi, A., Cotter, P., Kholodenko, B.N., 2015. DYVIPAC: an integrated analysis and visualisation framework to probe multidimensional biological networks. Scientific Reports 5, 12569., doi:10.1038/srep12569.

40. Nobis, C.C., Dubeau Laramée, G., Kervezee, L., Maurice De Sousa, D., Labrecque, N., Cermakian, N., 2019a. The circadian clock of cd8 t cells modulates their early response to vaccination and the rhythmicity of related signaling pathways. Proceedings of the National Academy of Sciences 116, 20077–20086. URL: https://www.pnas.org/content/pnas/116/40/20077.full.pdf, doi:10.1073/pnas.1905080116.

41. Nobis, C.C., Dubeau Laramée, G., Kervezee, L., Maurice De Sousa, D., Labrecque, N., Cermakian, N., 2019b. The circadian clock of CD8 T cells modulates their early response to vaccination and the rhythmicity of related signaling pathways. Proceedings of the National Academy of Sciences 116, 20077. URL: http://www.pnas.org/content/116/40/20077.abstract, doi:10.1073/pnas.1905080116.

42. Perley, J.P., Mikolajczak, J., Buzzard, G.T., Harrison, M.L., Rundell, A.E., 2014. Resolving early signaling events in T-cell activation leading to IL2 and FOXP3 transcription. Processes 2, 867–900. URL: https://www.mdpi.com/2227-9717/2/4/867.

43. Phillips, A.C., Gallagher, S., Carroll, D., Drayson, M., 2008. Preliminary evidence that morning vaccination is associated with an enhanced antibody response in men. Psychophysiology 45, 663–666.

44. Portaluppi, F., Tiseo, R., Smolensky, M.H., Hermida, R.C., Ayala, D.E., Fabbian, F., 2012. Circadian rhythms and cardiovascular health. Sleep Med Rev 16, 151–66. doi:10.1016/j.smrv.2011.04.003.

45. Sato, S., Sakurai, T., Ogasawara, J., Shirato, K., Ishibashi, Y., Oh-ishi, S., Imaizumi, K., Haga, S., Hitomi, Y., Izawa, T., et al., 2014. Direct and indirect suppression of interleukin-6 gene expression in murine macrophages by nuclear orphan receptor rev-erb*α*. The Scientific World Journal 2014.

46. Wang, C., Barnoud, C., Cenerenti, M., Sun, M., Caffa, I., Kizil, B., Bill, R., Liu, Y., Pick, R., Garnier, L., et al., 2023. Dendritic cells direct circadian anti-tumour immune responses. Nature 614, 136–143.

47. Wang, K., Wang, W., Liu, X., 2006. Viral infection model with periodic lytic immune response. Chaos, Solitons and Fractals 28, 90–99. URL: https://www.sciencedirect.com/science/article/pii/S0960077905004881, 10.1016/j.chaos.2005.05.003.

48. Wang, S., Lin, Y., Yuan, X., Li, F., Guo, L., Wu, B., 2018. Rev-erb*α* integrates colon clock with experimental colitis through regulation of nf*κ*b/nlrp3 axis. Nature communications 9, 1–12.

49. Young, M.R.I., Matthews, J.P., Kanabrocki, E.L., Sothern, R.B., RoitmanJohnson, B., Scheving, L.E., 1995. Circadian rhythmometry of serum Interleukin-2, Interleukin-10, Tumor Necrosis Factor-*α*, and GranulocyteMacrophage Colony-Stimulating factor in men. Chronobiology International 12, 19–27. doi:10.3109/07420529509064496.

50. Zheng, Y., Balakrishnan, V., Buzzard, G., Geahlen, R., Harrison, M., Rundell, A., 2005. Modeling and analysis of early events in T-lymphocyte antigen-activated intracellular-signaling pathways. Journal of Computational and Applied Mathematics 184, 320–341. URL: https://ui.adsabs.harvard.edu/abs/2005JCoAM.184..320Z.

